# CDH1 loss promotes diffuse-type gastric cancer tumorigenesis via epigenetic reprogramming and immune evasion

**DOI:** 10.1101/2023.03.23.533976

**Authors:** Gengyi Zou, Yuanjian Huang, Shengzhe Zhang, Kyung-Pil Ko, Bongjun Kim, Jie Zhang, Vishwa Venkatesan, Melissa P. Pizzi, Yibo Fan, Sohee Jun, Na Niu, Huamin Wang, Shumei Song, Jaffer A. Ajani, Jae-Il Park

**Author notes:** Correspondence: Jae-Il Park. ^†^These authors contributed equally.

## Abstract

This study investigates diffuse-type gastric adenocarcinoma (DGAC), a deadly and treatment-resistant cancer. It reveals that CDH1 inactivation occurs in a subset of DGAC patient tumors, leading to the identification of two distinct DGAC subtypes. The findings emphasize the importance of understanding DGAC’s molecular diversity for personalized medicine in patients with CDH1 inactivation.

## Abstract

Diffuse-type gastric adenocarcinoma (DGAC) is a deadly cancer often diagnosed late and resistant to treatment. While hereditary DGAC is linked to *CDH1* gene mutations, causing E-Cadherin loss, its role in sporadic DGAC is unclear. We discovered CDH1 inactivation in a subset of DGAC patient tumors. Analyzing single-cell transcriptomes in malignant ascites, we identified two DGAC subtypes: DGAC1 (CDH1 loss) and DGAC2 (lacking immune response). DGAC1 displayed distinct molecular signatures, activated DGAC-related pathways, and an abundance of exhausted T cells in ascites. Genetically engineered murine gastric organoids showed that *Cdh1* knock-out (KO), *Kras^G12D^*, *Trp53* KO (EKP) accelerates tumorigenesis with immune evasion compared to *Kras^G12D^*, *Trp53* KO (KP). We also identified EZH2 as a key mediator promoting CDH1 loss-associated DGAC tumorigenesis. These findings highlight DGAC’s molecular diversity and potential for personalized treatment in CDH1-inactivated patients.

## Introduction

Gastric adenocarcinoma (GAC) is the 4^th^ most common cause of cancer deaths worldwide (Sung et al., 2021). GAC is mainly divided into intestinal-type gastric adenocarcinoma (IGAC, 50%), diffuse-type gastric adenocarcinoma (DGAC, 30%), and mixed (Iyer et al., 2020). DGAC is histologically characterized by poor differentiation, loss of cell adhesion proteins, fibrosis, and infiltration. Unlike IGAC, DGAC is relatively more often observed in younger, female, and Hispanic population than in older, male, and non-Hispanic ones (Chen et al., 2016; Wang et al., 2020). While the incidence of IGAC has declined due to H. Pylori (HP) therapy and lifestyle improvements over the past few decades, the number of DGAC cases has remained constant or has risen (Henson et al., 2004; Assumpcao et al., 2020).

DGAC tends to metastasize to the peritoneal cavity, which makes it difficult to diagnose early by imaging. In addition, isolated tumor cells or small clusters of tumor cells infiltrate in unpredictable patterns. Thus, DGAC is often detected at a late stage, leading to a poor prognosis. For such patients, curative resection is not possible.

Systemic therapy is the main option for potentially prolonging survival and improving symptoms (Muro et al., 2019; Ajani et al., 2022). Despite the distinct features of DGAC in both a molecular basis and therapy resistance, the first-line treatment options are not specific for DGAC (Garcia-Pelaez et al., 2021; Ajani et al., 2022). Systemic therapy with targeted therapy has shown limited benefits (Selim et al., 2019; Korfer et al., 2021). In parallel, immune checkpoint inhibitors (ICIs) have been used recently. The advent of first-generation ICIs that target Cytotoxic T-Lymphocyte Antigen 4 (CTLA4) and Programmed death-ligand (PD-L1) has brought a paradigm shift in the treatment of various advanced cancers (Mazzarella et al., 2019). Nivolumab (PD-1 inhibitor) can be either combined with chemotherapy as first-line treatment or used as monotherapy as later-line treatment in Asia (Boku et al., 2021; Janjigian et al., 2021). Pembrolizumab (PD-1 inhibitor) showed a promising outcome treating GAC with high microsatellite instability or high tumor mutational burden (Wainberg et al., 2021). However, DGAC imposes major difficulty in the clinic and available therapies perform poorly. Generally, DGAC has immunosuppressed stroma and is genomically stable (Teng et al., 2015; Ge et al., 2018). Given the limited therapeutic options for DGAC, it is imperative to understand the biology of DGAC, which may establish a groundwork for developing new targeted therapies for DGAC. Furthermore, for maximizing therapeutic efficacy, it is crucial to identify patients who can most benefit from specific treatment options. Nevertheless, to date, DGAC patient stratification by molecular signatures has not been achieved.

Hereditary DGAC, as a minor proportion of DGAC (1–3%), is mainly characterized by germline mutations in the *CDH1* gene that encodes E-Cadherin (Blair et al., 2020). However, other than hereditary DGAC, the role of CDH1 loss in DGAC tumorigenesis is unclear. Cell-to-cell adhesion is a crucial phenomenon for maintaining tissue morphogenesis and homeostasis, as well as for regulating cell differentiation, survival, and migration. E-Cadherin mediates cell-to-cell adhesion, which is essential for determining the proliferation specificity and differentiation of epithelial cells and preventing invasion (van Roy and Berx, 2008). To understand the impact of *CDH1* loss on DGAC tumorigenesis, we analyzed single-cell transcriptomes of cryopreserved peritoneal carcinomatosis (PC) from 19 DGAC patients and identified two subtypes of DGACs exhibiting specific molecular signatures including E-Cadherin loss and immune landscape remodeling. To further verify our in-silico analysis, we generated and characterized a genetically engineered gastric organoid (GO) model that recapitulates E-Cadherin inactivation-associated DGAC tumorigenesis. This study stratifies DGAC patients by single-cell transcriptomics and elucidates the unexpected role of E-Cadherin loss in transcriptional reprogramming and immune evasion, providing novel insights into E-Cadherin loss-associated DGAC tumorigenesis.

## Results

### CDH1 inactivation in DGAC

To explore the role of CDH1 in DGAC tumorigenesis, we examined the genetic alterations and protein levels of CDH1 in DGAC. According to cBioPortal, 25% of tumor from the DGAC patients showed *CDH1* gene alterations, including mutations and deep deletions (**Fig. 1A**). We also assessed the CDH1 protein expression in the tissue microarray of 114 DGAC patients’ tumor samples (patient information was listed in **Table S4**). Immunohistochemistry (IHC) showed that 37.72% of DGAC patients were CDH1 negative, 37.72% exhibited low CDH1 expression, and 24.56% displayed high CDH1 expression (**Fig. 1B**), which was also quantified with histochemical scoring assessment (H-score) of each slide (**Fig. 1C**). Next, we determined the transcriptional signature of DGAC at the single-cell transcriptomics level by analyzing single-cell RNA-seq (scRNA-seq) datasets of PC cells from 19 stage IV DGAC patients (**Fig. 1D, Table S5**) (Wang et al., 2021). After data integration and normalization, a total of 30 cell clusters were generated according to distinctive gene expression patterns (**Fig. 1E**, **fig. S1A, B, Table S6**). We re-clustered the datasets as the mega clusters according to Leiden-based UMAP (**Fig. 1F**). To conduct the precise subtyping of DGAC, we reanalyzed the scRNA-seq datasets with only epithelial cells (**Fig. 1G**, **fig. S1C, Table S7**). An unsupervised pair-wise correlation analysis showed that the combined datasets of 19 DGAC patients were divided into two major subtypes (DGAC1 and DGAC2) (**Fig. 1H**). To comparatively analyze DGAC 1 and 2 according to their clinical information (**Table S5**), we have thoroughly examined the available data and compared various clinical and pathological features between the two subtypes. Upon analysis, we did not observe significant differences in survival time, race, gender, or age between DGAC1 and DGAC2 subtypes (**fig. S1D, S1E, S1G, and S1I**). Regarding pathological features, DGAC1 had a higher proportion (DGAC1: 3/11, 27.3%; DGAC2: 1/8, 12.5%) of patients with non-signet ring cell carcinoma (**fig. S1F**). A notable distinction in metastatic sites was displayed. DGAC1 patients exhibited a higher prevalence of metastatic sites compared to DGAC2. This observation suggests potential differences in the metastatic behavior of the two subtypes (**fig. S1H**). The transcriptional signature of DGAC1 epithelial cell clusters was distinct from that of DGAC2 (**Fig. 1I, J, and** **Table S8**). In line with the heterogeneity of CDH1’s genomic alterations and expression in DGAC patients (**Fig. 1A, B**), the DGAC1 subtype exhibited a relatively lower expression of *CDH1* compared to DGAC2 (**Fig. 1K, L**), indicating that the unsupervised pair-wise subtyping can also stratify DGAC patients by *CDH1* expression. While tissue microarray analysis showed that within the cohort of 114 DGAC patients, where 37.7% of patients exhibit low CDH1 levels (**Fig. 1B, C**), this subset of patients may have been classified into DGAC1 or DGAC2 based on the differential expression of other signature genes specific to each cluster, rather than solely relying on CDH1 expression. We also identified the molecular signatures of DGAC1 and DGAC2 (**Fig. 1M**). The gene list for calculating signature scores, including the DGAC1 and DGAC2 signatures, comprised the top 50 highly variable genes from each subgroup. (**Fig. 1M, Table S8**). These results identify two distinct subtypes of DGACs by distinct molecular signatures and CDH1 expression.

**Figure 1.**
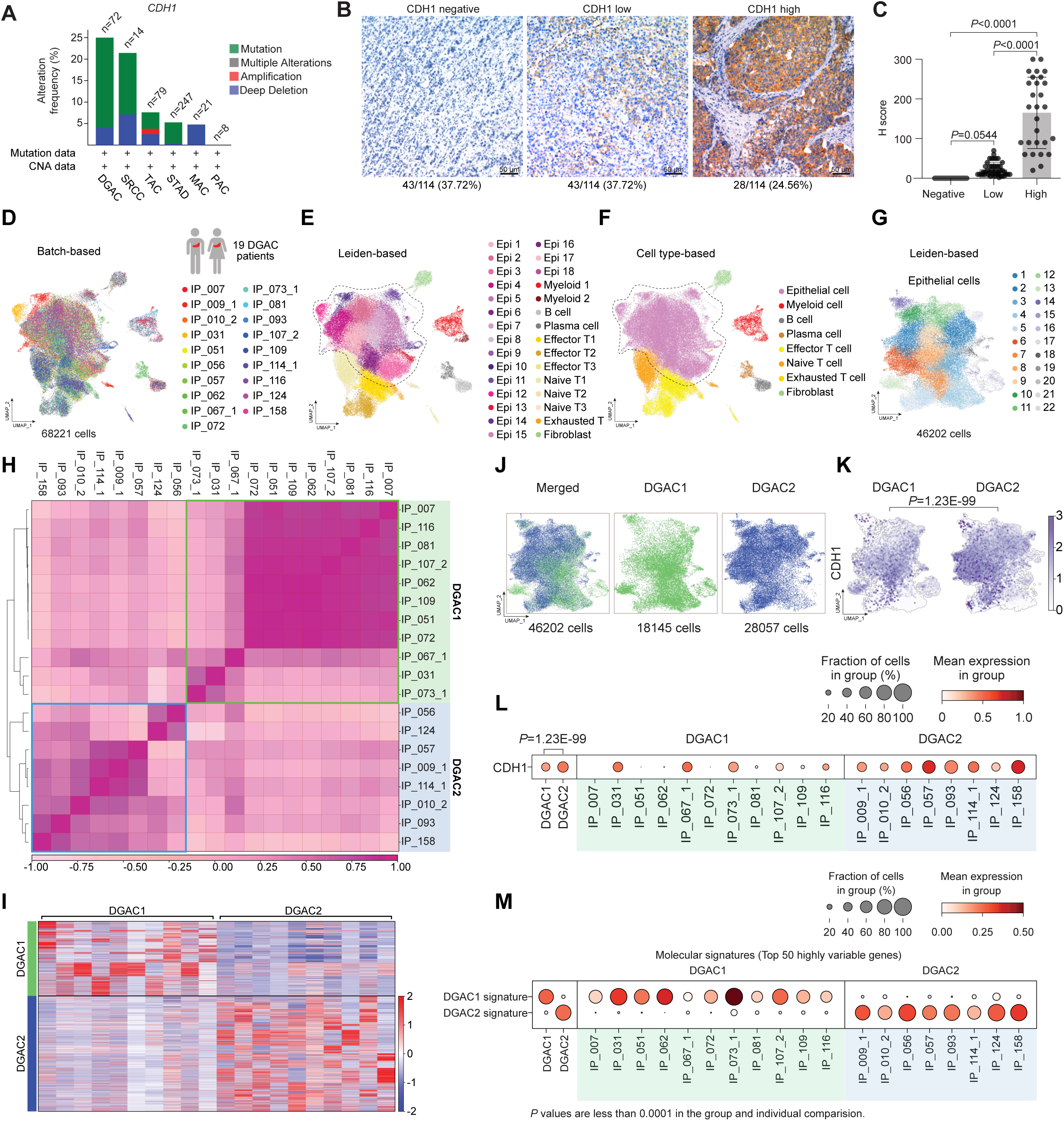
CDH1 inactivation in DGAC patient tumor cells. **A.** Genetic alteration of the *CDH1* based on the cBioPortal stomach cancer datasets (http://www.cbioportal.org). n represents the total patients number enrolled in each subtype. DGAC, diffuse-type gastric adenocarcinoma; SRCC, signet ring cell carcinoma; TAC, tubular adenocarcinoma; STAD, stomach adenocarcinoma; MAC, mucinous adenocarcinoma; PAC, papillary adenocarcinoma. **B, C.** IHC staining of CDH1 in 114 DGAC patient tumor samples. The representative images are shown (B). Quantification of H score of CDH1 expression (C). *P* values were calculated using the one-way ANOVA; error bars: standard deviation (SD). Clinical information of 114 DGAC patients was showed in Table S4. **D.** Merged batch-based integrated UMAPs of 19 DGAC patients; integration package: Harmony. Clinical information of 19 DGAC patients was showed in Table S5. **E.** Merged Leiden-based integrated UMAP of 19 DGAC patients. Dashed line circle: epithelial cells. Epi: epithelial cells; Myeloid: myeloid cells; Effector T: effector T cells; Naïve T: naïve T cells; Exhausted T: exhausted T cells. **F.** Merged cell type-based UMAP of 19 DGAC patients. All cells were re-clustered according to the Leiden clusters and gathered as mega clusters. Dashed line circle: epithelial cells. **G.** Epithelial cells were re-clustered by Leiden. **H.** Correlation matrix plot of epithelial cells showing pair-wise correlations among all samples above. The dendrogram shows the distance of each dataset based on principal component analysis, and the Pearson correlation is displayed with a color spectrum. Groups of patients were categorized by dendrogram and correlation. **I.** Type-based heatmap of epithelial cells of merged datasets in 19 DGAC patients. Top 100 highly variable genes of each type were showed in Table S8. **J.** Type-based integrated and separated UMAPs of DGAC1 and DGAC2. **K.** Feature plots of epithelial cells displaying *CDH1* expression. **L.** Dot plots of epithelial cells of *CDH1* expression in different DGAC groups and individual patients. **M.** Molecular signatures of DGAC1 and DGAC2 patients. Top 50 highly variable genes were used to calculate the molecular signature of each group. Gene list was showed in Table S8. Dot plots of epithelial cells of each molecular signature in different subtypes and individual patient.

## Molecular characterization of DGAC subtypes

Next, we characterized the molecular subtypes of DGAC. Given that E-Cadherin downregulation is commonly observed in epithelial tumors and is a hallmark of the epithelial to mesenchymal transition (EMT), we checked the EMT scores based on the established gene set (**Table S9**). DGAC1 showed a higher EMT score compared to DGAC2 (**fig. S1J**). Extensive genomic analyses of GAC have found that DGACs display distinct activation of signaling pathways different from IGACs (Ooki and Yamaguchi, 2022). By scRNA-seq-based signaling scoring, we observed that the FGFR2, HIPPO, PI3K/AKT/MTOR, and TGFBETA pathways were enriched in DGAC1, which corresponds to decreased CDH1 expression compared to DGAC2 (**fig. S1K, S1P, S1M, and S1R**). Additionally, we noted that FGFR1 is inversely correlated with CDH1 expression and enriched in DGAC1 (**fig. S1L**). Conversely, the RHOA and MAPK pathways were enriched in DGAC2 (**fig. S1N, S1O, and S1Q**). Additionally, we analyzed the copy number variation (CNV) of DGACs by using normal stomach samples as a reference. We combined 29 scRNA-seq datasets of normal stomach samples (Normal) with the previous 19 DGAC patients (Kim et al., 2022) (**fig. S2A**).

Except for the endothelial cell markers, the same marker panel was utilized as the previous DGAC subcategory process to annotate the cells into epithelial cells, myeloid cells, B cells, plasma cells, T cells, effector T cells, naïve T cells, exhausted T cells, fibroblasts, and endothelial cells (**fig. S1A, S2B, and S2C**). Leiden-based UMAP exhibited the same cell types as the DGAC stratification analysis (**fig. S2D, S2E, and Table S10**), except that the endothelial cell cluster appeared due to the normal tissue (**fig. S2C**). According to the previously identified DGAC subgroups, we separated the UMAP as Normal, DGAC1, and DGAC2 (**fig. S2F, fig. S2G**). Although the epithelial cells were defined as EPCAM^high^ clusters among all groups, epithelial cells from the Normal group were clearly isolated from the major epithelial cell population of the merged datasets (**fig. S2G**). CNV patterns were different between DGAC1 and DGAC2 (**fig. S2H**). We observed notable differences between DGAC1 and DGAC2 regarding copy number gains (GOF) on specific chromosomes. In DGAC1, we observed more pronounced copy number gains on chromosomes 3, 9, 19, and X, while in DGAC2, there were increased copy number gains on chromosomes 1, 8, 11, 17, 20, and 21. These differences in copy number alterations were found to be statistically significant, as indicated by the adjusted *P* values (**fig. S2I, S2J**). These results indicate the heterogeneity of DGAC with differentially activated signaling pathways.

## Immune landscape remodeling with T cell exhaustion in DGAC1

Having determined the molecular signatures of DGAC tumor cells, we next analyzed immunological response associated with DGAC ascites. Intriguingly, scRNA-seq-based immune cell profiling showed that compared to DGAC2 where immune cells barely existed, DGAC1 was highly enriched with immune cells, including T cells, B cells, and myeloid cells (**Fig. 2A-C**, **fig. S2K-L**). Additionally, we examined cellular networks among all cell clusters (DGAC1 vs. DGAC2) using a CellChat package that infers cell-to-cell functional interaction based on ligand-receptor expression (Jin et al., 2021).

**Figure 2.**
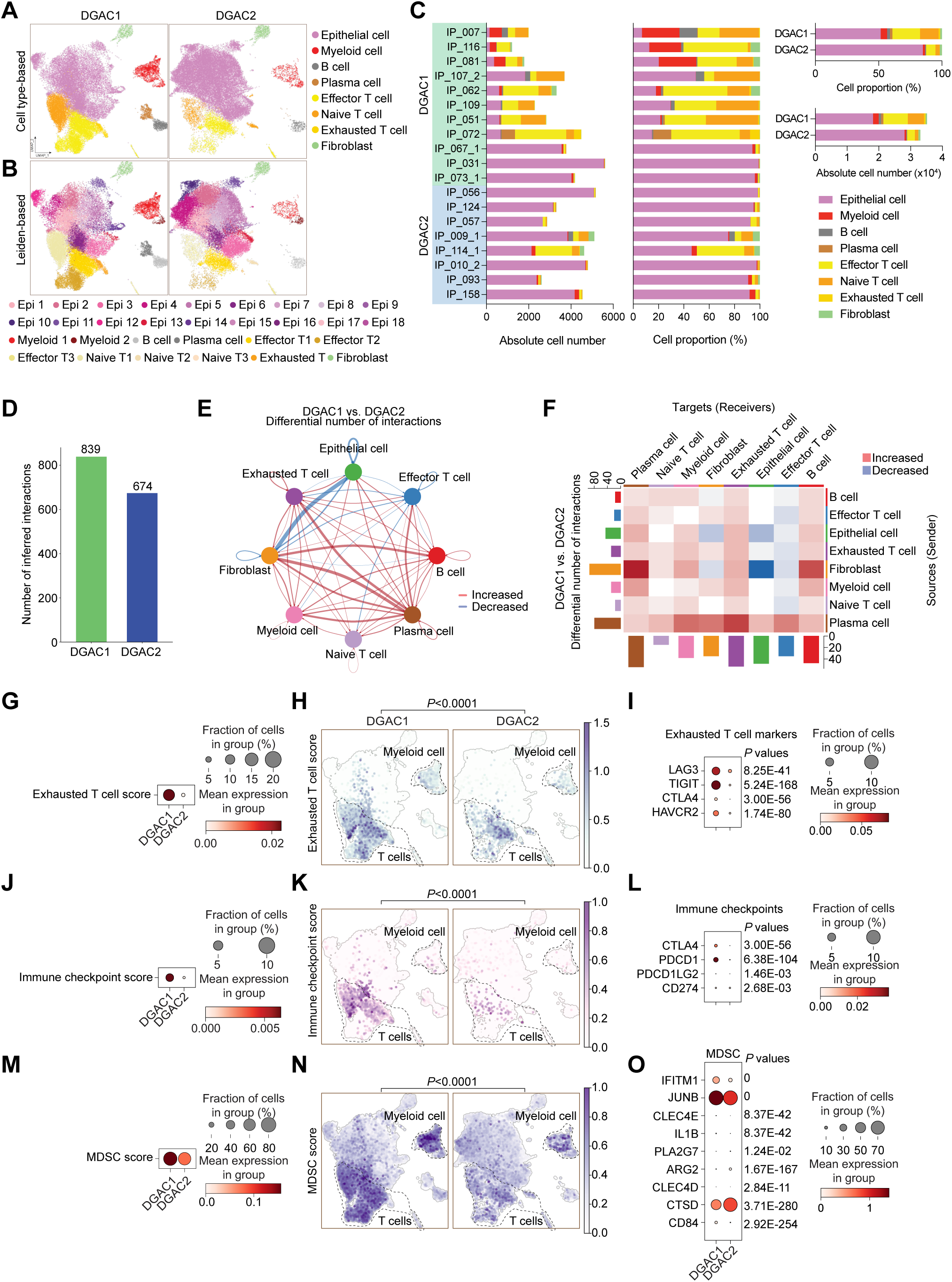
Comparative analyses of immune landscapes of DGAC subtypes A-B. Cell type-based and Leiden-based UMAPs of DGAC1 and DGAC2. **C.** Absolute and relative cell proportions of individual patients and DGAC subtypes. Patients list was ranked by the DGAC group that they belong. **D.** Total cell-cell interactions from DGAC1 and DGAC2 were analyzed by using the CellChat package. More interactions were found in DGAC1. **E, F.** Differential number of interactions between DGAC1 and DGAC2 using circle plots (E) and heatmap (F). Red (or blue) colored edges (E) and squares (F) represent increased (or decreased) signaling in the DGAC1 compared to DGAC2. **G-I.** Score-based dot plot (G), feature plots (H), and dot plot of individual marker gene (I) of exhausted T cell score (markers are included in that score: *LAG3*, *TIGIT*, *CTLA4*, and *HAVCR2*). Genes that included in score analysis were showed in Table S9. *P* values were calculated by using a *t*-test. **J-L.** Score-based dot plot (J), feature plots (K), and dot plot of individual marker gene (L) of immune checkpoint score (markers are included in that score: *CTLA4*, *PDCD1*, *PDCD1LG2*, and *CD274*). Genes that included in score analysis were showed in Table S9. *P* values were calculated by using a *t*-test. **M-O.** Score-based dot plot (M), feature plots (N), and dot plot of individual marker gene (O) of exhausted T cell score (markers are included in that score: *IFITM1*, *JUNB*, *CLEC4E*, *IL1B*, *PLA2G7*, *ARG2*, *CLEC4D*, *CTSD*, and *CD84*). Genes that included in score analysis were showed in Table S9. *P* values were calculated by using a *t*-test.

Compared to DGAC2, DGAC1 showed relatively more inferred interactions among different cell types (**Fig. 2D**). According to the differential number of interactions, the interactions between fibroblast and epithelial and endothelial cells were decreased, while widespread increased interactions were found in DGAC1 compared to DGAC2 (**Fig. 2E**). Notably, exhausted T cells, as a receiver, showed the most increased interactions compared with other T cells in DGAC1, which is the major population among all immune cells (**Fig. 2F**). Gene Set Enrichment Analysis (GSEA) identified the pathways that are enriched in DGAC1 with six gene sets, including Gene sets derived from the Gene Ontology Biological Process (GOBP), and five canonical pathways gene sets (REACTOME, WP, BIOCARTA, PID, and KEGG) (**fig. S3A-F, Table S11-S16**). Except for REACTOME (**fig. S3B**), T cell-related immune response pathways were enriched in DGAC1 based on the other five gene sets (**fig. S3A, S3C, S3D-F**). Consistent with the CellChat prediction and GSEA results, DGAC1 showed the significant upregulation of T cell exhaustion markers (LAG3, TIGIT, CTLA4, and HAVCR2) and the increased T cell exhaustion score, compared to DGAC2 (**Fig. 2G-I**). Similarly, immune checkpoints-related genes (CTLA4, PDCD1, PDCD1LG2, and CD274) and their score were markedly upregulated in DGAC1 over DGAC2 (**Fig. 2J-L**). In addition to T cell analysis, we also examined myeloid-derived suppressor cells (MDSC) and macrophage polarization. MDSC score is relatively higher in DGAC1 than DGAC2 (**Fig. 2M-O**). Meanwhile, most of M1 and M2 macrophage polarization maker expression is enriched in DGAC1 compared to DGAC2 (**fig. S3G-H**). These results suggest that compared to DGAC2, the DGAC1 subtype exhibits distinct immune remodeling featured by T cell exhaustion and increased expression of the genes associated with immune checkpoints.

### Cdh1 loss induces neoplasia in conjunction with *Trp53* KO and *Kras^G12D^*

To validate the in silico results, we utilized murine GOs that enable multiple genetic engineering with immediate phenotype analyses. *Cdh1* deficiency results in early-stage DGAC phenotype in a mouse model (Mimata et al., 2011; Hayakawa et al., 2015). Nevertheless, other genes need to be included to recapitulate DGAC tumorigenesis. The genes encoding the receptor tyrosine kinase (RTK)-RAS signaling pathway and the *TP53* gene were profoundly disrupted in DGAC (Cancer Genome Atlas Research, 2014; Cristescu et al., 2015). *KRAS* and *TP53* were genetically altered in 13.19% and 36.11% of DGAC cases, respectively, as per cBioPortal analysis (**Fig. 3A**). Furthermore, we observed that the PI3K/AKT/mTOR pathway is activated in DGAC1 (**fig. S1M**). Therefore, to create preneoplastic or neoplastic conditions to determine the impact of CDH1 loss on DGAC tumorigenesis (Till et al., 2017), we genetically manipulated three genes (*Cdh1*, *Trp53*, and *Kras*) in GOs. Briefly, from the *Cdh1* wild type (WT) and *Kras ^LSL-G12D/+^*; *Trp53^fl/fl^*mice, gastric epithelial cells were isolated to culture them into GOs (**Fig. 3B**). Subsequently, using the Cre-LoxP recombination and CRISPR-based genetic manipulation, we established two lines of GOs carrying *Kras^G12D/+^*and *Trp53* deletion in combination with *Cdh1* KO (KP: *<u>K</u>ras^G12D/+^*; *Tr<u>p</u>53* KO [KP], *Cdh1/<u>E</u>-Cadherin* KO; *<u>K</u>ras^G12D/+^*; *Tr<u>p</u>53* KO [EKP]) (**Fig. 3B**). Genetic modifications were validated by PCR-based genotyping and genomic DNA sequencing and immunofluorescence (IF) staining (**fig. S4A-C**, **Fig. 3G**). Meanwhile, we monitored their sizes and numbers by macroscopic analyses during passages to maintain the stable culture process during passages (**Fig. 3C, D**). Unlike WT GOs growing as a single layer of epithelial cells, KP and EKP GOs displayed multilayered epithelium (**Fig. 3E**). Notably, compared to WT and KP, EKP GOs exhibited abnormal morphology such as vacuolization and cell adhesion loss along with cell hyperplasia (**Fig. 3E**). Additionally, EKP GOs were hyperproliferative compared to WT and KP GOs, assessed by immunostaining of MKI67, a cell proliferation marker (**Fig. 3F, H**).

**Figure 3.**
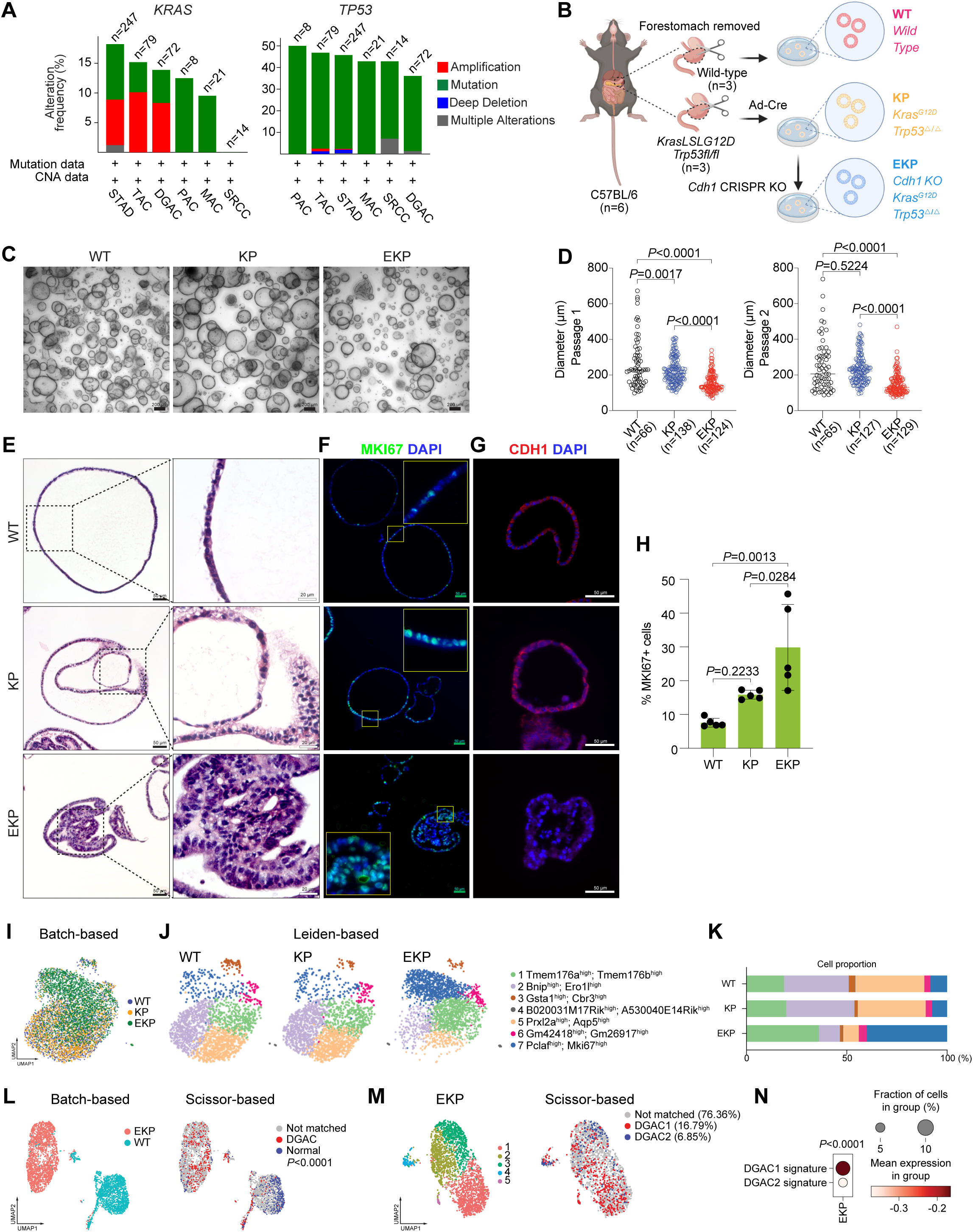
Establishment of genetically engineered gastric organoids with CDH1-inactivation. **A.** Genetic alteration of the *KRAS*, and *TP53* genes based on the cBioPortal. n represents the total patients number enrolled in each subtype. DGAC, diffuse-type gastric adenocarcinoma; SRCC, signet ring cell carcinoma; TAC, tubular adenocarcinoma; STAD, stomach adenocarcinoma; MAC, mucinous adenocarcinoma; PAC, papillary adenocarcinoma. **B.** Illustration of the workflow for stomach tissue collection and dissociation, gene manipulation of the gastric organoids (GOs), GOs culture, and representative image of GOs. Three GO lines were generated, including WT, KP, and EKP. WT mice and KP mice were sacrificed to collect stomach tissue. After removing forestomach, stomach tissue was dissociated into single cell and culture as organoids. Adeno-Cre virus was used to treat *Kras^LSL-G12D^; Trp53^fl/fl^* organoids to generate KP organoids, followed by nutlin-3 selection. After selection, EKP organoids were generated using CRISPR-mediated *Cdh1* KO from KP GOs. **C.** Representative images of WT, KP, and EKP GOs at passage day 8. Scale bar: 200 μm. **D.** Growth analysis for WT, KP, and EKP GOs in two passages at day 8 of each passage. P values were calculated using the one-way ANOVA; error bars: SD. Numbers below each label represent the number of organoids. **E.** Hematoxylin and eosin (H & E) staining of WT, KP, and EKP GOs. **F.** MKI67 staining of WT, KP, and EKP GOs (n=5). **G.** CDH1 staining of WT, KP, and EKP GOs (n=5). **H.** Statistics analysis of MKI67 staining (Figure 3F). *P* values were calculated using the one-way ANOVA; error bars: SD. The representative images are shown. **I.** Batch-based UMAPs of WT, KP, and EKP GOs. The Harmony integration package was used to remove the batch effect. **J.** Leiden-based clustering UMAPs of WT, KP, and EKP GOs. Cell clusters were named by the most highly variable genes. **K.** Cell proportion analysis of WT, KP, and EKP GOs. Each color represents a different cell type. The color code is based on the cell types shown in Figure 3J. **L.** Batch-based and Scissor-based UMAP of WT and EKP GOs generated by Scissor package. TCGA datasets of normal stomach and DGAC patients were utilized. **M.** Cluster-based and Scissor-based UMAP of EKP GOs generated by Scissor package. DGAC1 and DGAC2 datasets were utilized to perform the comparison. **N.** Dot plots of EKP GOs of DGAC1 and DGAC2 molecular signatures. Top 50 highly variable genes were used to calculate the molecular signature of each DGAC subtype. Gene list was showed in Table S8.

We next interrogated the mechanism of Cdh1 loss-associated DGAC tumorigenesis by performing multiplex scRNA-seq of WT, KP, and EKP GOs (**fig. S4D**). Each group was tagged with two Cell Multiplexing Oligo (CMO) tags, then pooled together with the same number of cells after being counted. All datasets were integrated with the Harmony algorithm (Korsunsky et al., 2019) to minimize the batch effect (**fig. S4E**). WT, KP, and EKP GOs were merged well in a batch-based UMAP (**Fig. 3I**). To identify the gene signature of each cell cluster, we generated a heatmap to calculate the top 5,000 highly variable genes (**fig. S4F**). Each UMAP and heatmap represented the different cell distribution among three types of GOs (**Fig. 3J, K, fig. S4G-I, and Table S17**). Notably, Aquaporin 5 (Aqp5), a gastric tissue stem cell marker (Tan et al., 2020), was decreased in EKP compared to WT and KP (**Fig. 3K**).

To determine the pathological relevance of EKP GOs with human DGAC, we utilized a single-cell inferred site-specific omics resource (Scissor) analysis (Sun et al., 2022) and assessed the transcriptomic similarity between of EKP GOs and the bulk RNA-seq data of patients diagnosed with DGAC from the TCGA database. While using WT organoids as a reference and comparing the transcriptional signature, we observed that EKP organoids displayed similarities in gene expression features of human DGAC (**Fig. 3L**). To determine the subtype similarity, we compared the EKP scRNA-seq data with our own datasets (DGAC1 and DGAC2) rather than relying solely on the TCGA database. The analysis revealed that EKP organoids exhibited a greater resemblance to DGAC1 transcriptional signature compared to DGAC2 (**Fig. 3M**). Next, by comparing the expression levels of DGAC1 and DGAC2 signatures in EKP (**Fig. 1M**), we observed a higher presence of DGAC1 signature compared to DGAC2 (**Fig. 3N**). These data suggest that CDH1 loss combined with TP53 inactivation and KRAS hyperactivation (EKP) is sufficient to induce transformation and EKP organoids display similar transcriptional features to DGAC1, indicating pathological relevance of EKP GOs to human DGAC.

## *Cdh1* KO induces immune evasion of tumor cells

Having determined distinct immune remodeling with T cell exhaustion in the DGAC1 subtype where CDH1 is downregulated (**Fig. 2**), we asked whether genetic ablation of *CDH1* contributes to immune evasion of DGAC. To test this, we established KP and EKP GO-derived cell lines in 2D culture with minimum growth factors (culture medium: DMEM Complete Medium with 10% fetal bovine serum) for allograft transplantation (**Fig. 4A**). Unlike WT GOs that failed to grow in 2D culture, both KP and EKP cells grew in 2D culture and were maintained well at multiple passages. Then, KP and EKP cell lines derived from C57BL/6 strain were used for transplantation into C57BL/6 mice. The morphological characteristics of KP and EKP cells exhibited notable differences. KP cells exhibited a compact and tightly packed phenotype, forming densely clustered colonies, while EKP cells displayed a more loosely-arranged and dispersed morphology, lacking the cohesive structure of KP cells (**Fig. 4A**). Of note, there was no significant difference in cell proliferation between KP and EKP cells (**Fig. 4B**). However, transplantation results showed that tumor incidence and volume of EKP tumors was markedly higher than KP tumors (tumor incidence rates: EKP [91.7%] vs. KP [16.7%]) (**Fig. 4C-E**). Histologically, EKP tumors exhibited poorly differentiated tumor cells, the feature of DGAC (**Fig. 4F**) with CDH1 loss (**Fig. 4H**) and increased cell proliferation (**Fig. 4G, I**).

**Figure 4.**
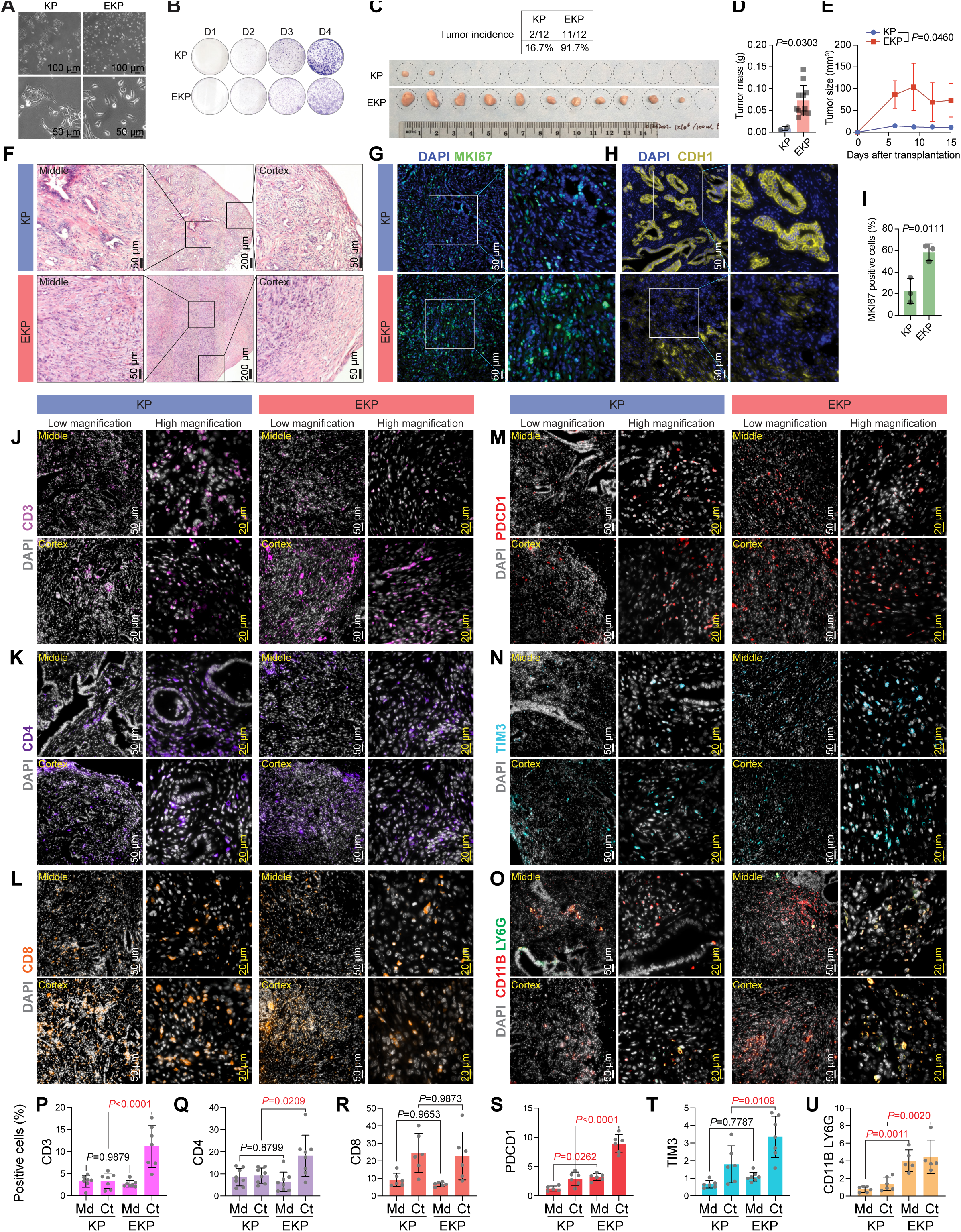
*CDH1* KO promotes KP-driven gastric tumorigenesis. **A.** Bright-field images of KP and EKP cells in low and high magnification. **B.** Crystal violet staining of KP and EKP GOs-derived cells. **C.** Bright-field images of KP and EKP allograft tumors; tumor incidence of allograft tumors. **D, E.** Plot for tumor mass (D) and tumor size (E) assessment of KP and EKP allografts. **F.** H & E staining of KP and EKP allograft tumors (n≥3). **G, H.** MKI67 (G) and E-Cadherin (H) staining of KP and EKP allograft tumors (n≥3). Left images: low magnification. Right images: high magnification. Scale bars were shown on the representative images. **I.** Statistics analysis of MKi67 staining in Figure 4G. *P* values were calculated using Student’s *t*-test; error bars: SD. **J-O.** CD3 (J), CD4 (K), CD8 (L), PDCD1 (M), TIM3 (N) staining and CD11B/LY6G co-staining (O) of KP and EKP allograft tumors (n≥3). Middle and Cortex represents the middle and cortex of the tumor, respectively. In each panel, left images showed low magnification, and right images showed high magnification. Scale bars were shown on the representative images. **P-U.** Statistics analysis of CD3 (P), CD4 (Q), CD8 (R), PDCD1 (S), TIM3 (T) staining and CD11B/LY6G co-staining (U). The positive cell percentage indicates the area of cells expressing a specific marker divided by the total field occupied cells stained by DAPI in the same area, which allows for normalization. Md: Middle; Ct: Cortex. *P* values were calculated using the one-way ANOVA; error bars: SD.

To further determine the impact of CDH1 loss on immune evasion, we performed the immunostaining of KP and EKP tumors. We observed CD3 (a marker for all T cells), CD4 (a marker for helper T cells), and TIM3 (a marker for exhausted T cells) are enriched in EKP tumor cortex compared with KP cortex (**Fig. 4J, 4K, 4N, 4P, 4Q, and 4T**), and the CD8 (a marker for killer T cells) expression is similar between KP and EKP tumors (**Fig. 4L and 4R**). PDCD1, a marker for exhausted T cells, showed increased expression in the middle and cortex of EKP compared with the same part of KP tumors (**Fig. 4M and 4S**). Furthermore, we performed LY6G (a marker for MDSCs) and CD11B (a marker for myeloid cells) co-staining on tumor slides of KP and EKP and observed the relatively higher enrichment of MDSC markers in EKP (**Fig. 4O and 4U**). These results suggest that CDH1 is a gatekeeper restricting the immune evasion of DGAC, confirming immune landscape remodeling associated with the DGAC1 subtype where CDH1 is inactivated.

## *Cdh1* depletion-activated EZH2 regulon promotes gastric tumorigenesis

Since CDH1 loss is associated with distinct molecular signatures of DGAC1 (**Fig. 1M**), we sought to identify key transcriptional regulatory modules (regulons) activated by *Cdh1* depletion. We integrated the scRNA-seq datasets of WT, KP, and EKP into batch-based and regulon pattern-based UMAPs (**Fig. 5A**). In the regulon activity-based UMAP, six major transcriptional clusters (0∼5) were identified (**Fig. 5A**). With the separated UMAP, we observed that WT and KP shared somewhat similar transcriptional landscape. However, EKP exhibited distinct features with an increased cluster 5 (**Fig. 5B**). To pinpoint essential regulons, we created an unbiased workflow (**Fig. 5C**). Based on the Z score of each regulon, we identified 32 regulons specific to EKP transcriptional profile, compared to those of WT and KO (**Fig. 5D**). Additionally, regulon specificity score (RSS) analysis showed the top 20 regulons specific to EKP (**Fig. 5E**). RSS-based top 20 regulons belonged to Z score-based regulons (**Fig. 5F, Table S18**). Both RSS and Z-score were used to quantify the activity of a gene or set of genes. Z-score was used to quantify the level of gene expression in a particular sample, while RSS was used to quantify the specificity of a gene set to a particular regulatory network or module (Kelley et al., 2016). According to TCGA-based upregulation in DGAC patients compared to normal stomach tissues, 13 regulons (Brca1, E2f1, E2f3, E2f7, E2f8, Ezh2, Gabpa, Gtf2b, Gtf2f1, Hmga2, Pole4, Sox4, and Tfdp1) were selected (**fig. S5A**). Next, we examined the regulons’ expression in organoids datasets.

**Figure 5.**
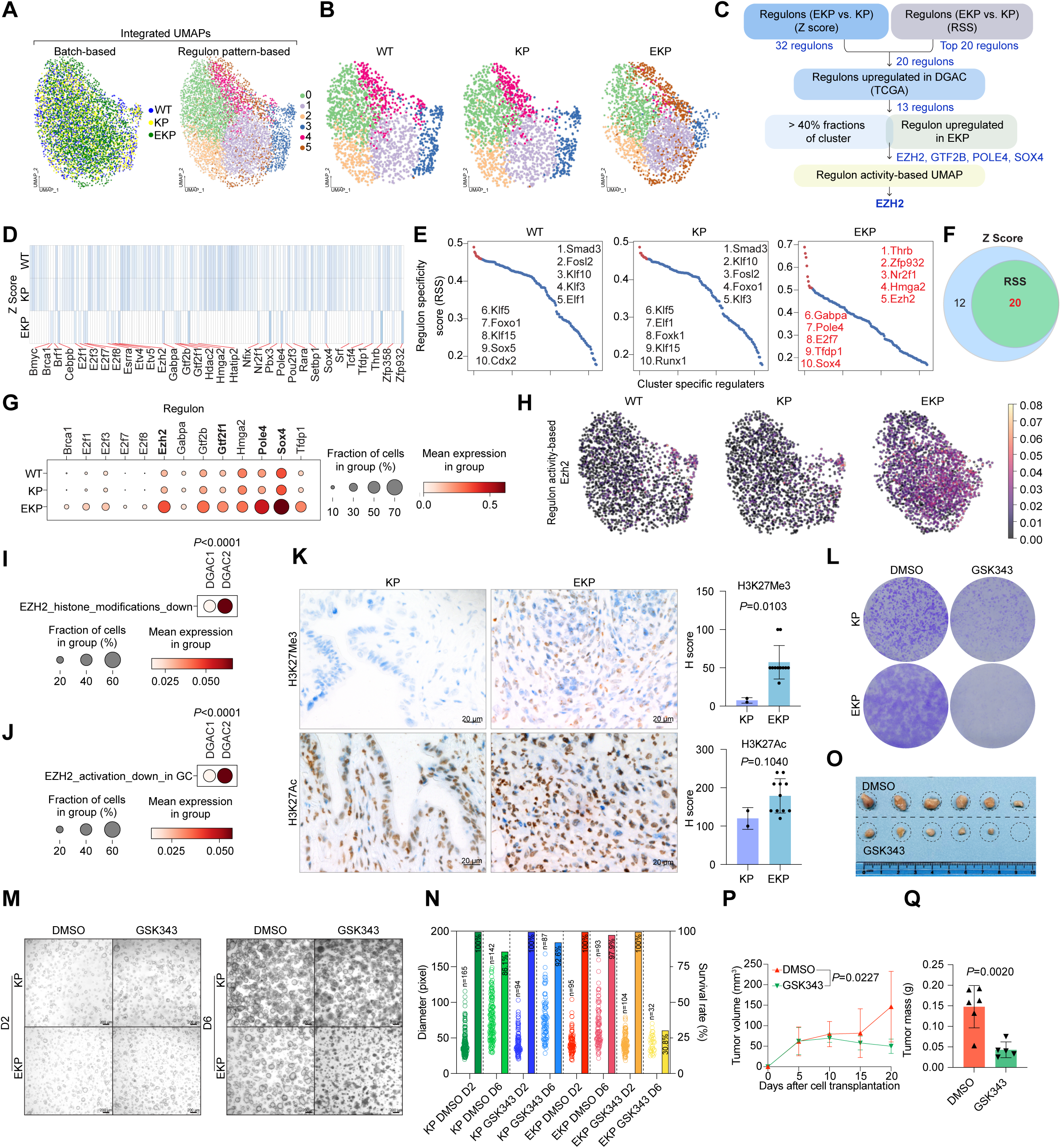
*CDH1* KO-activated EZH2 promotes gastric tumorigenesis. **A.** Integrated batch-based and regulon pattern-based UMAP for WT, KP, and EKP GOs. Six transcriptional modules were identified. **B.** Separated regulon patterns based UMAP for WT, KP, and EKP GOs. **C.** Flow chart of regulons selection process. **D.** Regulons enriched in WT, KP, and EKP GOs, based on Z Score. 32 regulons were highly expressed in EKP samples compared to WT and KP. **E.** Regulons enriched in WT, KP, and EKP GOs, based on Regulon Specificity Score (RSS). The top 20 were selected by Z score. The whole regulon list based on RSS was showed in Table S18. **F.** Venn diagram for the regulons from figure 5D and 5E. 20 regulons were overlapped. **G.** Dot plot of the regulons (WT, KP and EKP GOs) increased in TCGA DGAC patients. **H.** Regulon activity-based UMAP of Ezh2 in WT, KP, and EKP GOs. The cells with lighter color represent regulated by Ezh2. **I, J.** Dot plots of EZH2 downstream target genes (I, genes which are downregulated by EZH2 activation through histone modification; J, genes which are downregulated by EZH2 activation reported in gastric cancer) scores in the epithelial cells of DGAC1 and DGAC2. Gene list of EZH2 targeted genes was listed in Table S9. **K.** The level of H3K27Ac and H3K27Me3 expression in KP and EKP allografts. Quantification was displayed. **L.** Crystal violet staining of KP and EKP cells after GSK343 (EZH2 inhibitor, 10 μM, 96 hrs). **M.** Bright field images (M) of KP and EKP GOs after treating with GSK343 (EZH2 inhibitor, 10 μM, 96 hrs). D2: day2; D6: day6. **N.** Statistical analysis of KP and EKP gastric organoid size and number in response to GSK343 treatment. The number of organoids (right Y-axis) and their size (left Y-axis) were assessed following treatment with GSK343. At Day 2 (D2), the number of organoids was determined for the image depicted in figure 5M, and this count was considered as 100% (n numbers are presented in the bubble plot). At Day 6 (D6), the number of organoids in the same field for each group was counted (n numbers also displayed in the bubble plot). The percentage of each group on D6 was calculated by dividing the number of viable organoids at D6 by the number at D2. The viable percentage is presented in the bar graph. **O-Q.** Transplantation of EKP cells followed by EZH2 inhibition. Bright-field images of EKP allograft tumors treated with DMSO and GSK343 (20 mg/kg) separately (O). Tumor growth curve of EKP allografts treated with DMSO and GSK343 (20 mg/kg) after cell subcutaneous transplantation (P). Tumor mass of EKP allografts treated with DMSO and GSK343 (20 mg/kg) after mice scarification (Q). *P* values were calculated using Student’s *t*-test; error bars: SD.

Compared to WT and KP, the expression of Ezh2, Gtf2b, Pole4, and Sox4 was obviously increased in EKP GOs with over 40% fractions of clusters (**Fig. 5G**). According to the regulon activity-based UMAP, Ezh2 displayed the highest score in EKP compared to WT and KP GOs (**Fig. 5H**, **fig. S5B**). To assess the pathological relevance of EZH2 to DGAC, we analyzed the expression of downstream target genes of EZH2 in the DGAC datasets (**Table S9**) (Yu et al., 2023). One gene list included the genes which are downregulated by EZH2 activation through histone modification (EZH2_histone_modification_down) (**Fig. 5I**, **fig. S5C, Table S9**); the other gene list included the genes which are downregulated by EZH2 activation that reported in gastric cancer (EZH2_activation_down_in_GC) (**Fig. 5J**, **fig. S5D, Table S9**). Compared to DGAC2 (CDH1 high), the EZH2_histone_modification_down and EZH2_activation_down_in_GC scores were relatively lower in DGAC1 (CDH1 loss) (**Fig. 5I, J**). EZH2 is a histone methyltransferase catalyzing the methylation of histone H3 lysine 27 (H3K27) to generate H3K27me3, which is associated with gene repression (Lee et al., 2007). Consistent with EZH2 regulon activation by *Cdh1* KO, H3K27Me3 was also increased in EKP tumors compared to KP, while no significant difference in H3K27Ac expression (**Fig. 5K**). Next, we treated EKP cells with GSK343, a specific inhibitor of EZH2 methyltransferase (Verma et al., 2012). EKP cells were more sensitive to GSK343 than KP in cell proliferation in vitro (**Fig. 5L**). Meanwhile, we conducted experiments to evaluate the effect of GSK343 on KP and EKP organoids (**Fig. 5M**). We observed that the number of EKP GOs was significantly decreased (30.8%) after GSK343 treatment, while the number of KP GOs was marginally affected by GSK343 (92.6%) of the organoids initially seeded (**Fig. 5N**). Additionally, allograft transplantation experiments showed the growth inhibitory effect of GSK343 on EKP tumorigenesis (**Fig. 5O, P, Q**). These results identify EZH2 as a key regulon contributing to tumorigenesis of CDH1 inactivation-associated DGAC.

## Discussion

The impact of CDH1 loss on sporadic DGAC tumorigenesis remains unknown. Single-cell transcriptomics-based unsupervised clustering identified two subtypes of DGAC: DGAC1 (CDH1-negative or downregulated) and DGAC2 (CDH1-positive). Unlike DGAC2 lacking ascites tumor cells-associated immunologic response, the DGAC1 subtype is enriched with exhausted T cells. Single-cell transcriptomics and transplantation assays showed that *Cdh1* KO, in conjunction with *Trp53* KO and *Kras^G12D^*, induces accelerated tumorigenesis and immune evasion. Moreover, EZH2 regulon specifically activated by CDH1 loss promotes DGAC tumorigenesis.

Patient stratification is crucial for improving therapeutic efficacy. Despite several studies classifying GAC patients (Ge et al., 2018; Fukamachi et al., 2019; Tong et al., 2019; Kim et al., 2020; Wang et al., 2020), such subtyping did not consider single-cell level cellular convolution, which might be insufficient to represent the full spectrum of DGAC features. Our stratification approach was based on the high dimensional transcriptional signatures at the single-cell level, immune cell profiling, and cellular network, which may complement limitations from the bulk analyses and likely better stratify DGAC patients. Indeed, our unsupervised subtyping by tumor cell transcriptome well matched with distinct immune cell properties (**Fig. 2A-C**). Furthermore, the application of CellChat and GSEA analysis led to the identification of T cell-related immune profiling as the dominant feature in DGAC1 (**Fig. 2D-F**, **fig. S3A-F**).

Interestingly, T cell exhaustion and immune checkpoint-related genes were notably enriched in DGAC1 compared to DGAC2 (**Fig. 2G-L**), confirmed by the transplantation experiments (**Fig. 4**). These results strongly suggest that DGAC1 patients might benefit from T cell-based ICIs, whereas DGAC2 patients might be ICI non-responders (**Fig. 2**).

Understanding the biology of cancer immune evasion is also imperative for improving cancer treatment. To date, how DGAC tumor cells evade immune surveillance remains elusive. Our transplantation assays showed that in conjunction with *Trp53* KO and *Kras^G12D^*, CDH1 loss is sufficient for immune evasion of DGAC (**Fig. 4**). In line with this, EKP allografts displayed increased expression of CD3, CD4, PDCD1, TIM3, and LY6G (**Fig. 4J-U**), also identified as molecular signatures of DGAC1 (**Fig. 2G-O**). These tantalizing results suggest a new role of CDH1 in restricting the immune evasion of tumor cells beyond its canonical role in cell-cell adhesion.

Tumors are immunogenically categorized into ‘hot’, ‘altered-excluded’, ‘altered-immunosuppressed’ and ‘cold’ (Galon and Bruni, 2019). The terms ’hot’ and ’cold’ describe T cell-infiltrated inflamed tumors and non-infiltrated tumors, respectively (Galon et al., 2006). Altered-immunosuppressed tumors have few CD8+ T cells, mainly at the tumor’s periphery, with immune-suppressing cells like MDSCs and regulatory T cells. In altered-excluded immune tumors, CD8+ T cells are absent, and the tumor microenvironment is dense and hypoxic, hindering immune cell survival (Galon and Bruni, 2019). Cold tumors, altered–immunosuppressed, or immune–excluded tumors, respond less favorably to ICIs and generally have a poorer prognosis compared to hot tumors, which tend to respond well to ICIs (Galon and Bruni, 2019; Lee and Ruppin, 2019). According to the immune profiling of EKP tumors (**Fig. 4**), which mimic DGAC1, it is highly probable that DGAC1 may correspond to hot or altered–immunosuppressed mixed tumors, while DGAC2 is likely to be classified as either cold tumors or altered– excluded immune tumors. Emerging evidence suggests that CDH1 loss may be associated with an inflamed phenotype (Stodden et al., 2015; Kaneta et al., 2020). E-Cadherin encoded by *CDH1* is an adhesion molecule responsible for maintaining cell-cell interactions and tissue integrity. Loss of CDH1 disrupts adherens junctions of tumor cells and subsequently disorganizes tumor architecture (Bruner and Derksen, 2018), which likely promotes immune cell infiltration.

Previously, two distinct molecular subtypes of GAC were introduced: mesenchymal phenotype (MP) and epithelial phenotype (EP) (Oh et al., 2018; Wang et al., 2020). Since its association with CDH1 downregulation and EMT (**fig. S1J**), the DGAC1 subtype might belong to the MP subtype, which displays poor survival and chemotherapy resistance (Oh et al., 2018). Unlike DGAC1, DGAC2 does not show CDH1 loss and EMT. DGAC exhibits frequent mutations in the *TP53, CDH1, RHOA, APC, CTNNB1, ARID1A, KMT2C, and PIK3CA* genes (Cancer Genome Atlas Research, 2014; Kakiuchi et al., 2014; Oliveira et al., 2015; Cho et al., 2019). Among these genes, *CDH1* and *RHOA* mutations are mainly observed in DGAC and not found in IGAC (Cancer Genome Atlas Research, 2014; Kakiuchi et al., 2014; Cho et al., 2019). As CDH1 and RHOA both play a crucial role in modulating the cytoskeleton, cell morphology, and cell migration (Handschuh et al., 1999; McBeath et al., 2004; O’Connor and Chen, 2013; Al-Ahmadie et al., 2016), the general histological features of DGAC are likely attributed to mutations in these genes, *CDH1* and *RHOA* (Ooki and Yamaguchi, 2022). DGAC2 displays high *CDH1* expression and no EMT gene expression (**Fig. 1J-1L**, **fig. S1J**). Additionally, DGAC2 shows RHOA signaling activation (**fig. S1N**). Thus, epithelial cell polarity loss and the diffuse-type cell (morphological) phenotype in DGAC2 might be due to *RHOA* mutations (Y42C) or RHOA signaling activation, whereas CDH1 loss is linked with DGAC1.

E-Cadherin mediates cell-cell interaction via homophilic interaction with other E-Cadherin proteins from neighboring cells. The cytoplasmic domain of E-Cadherin is physically associated with Catenin proteins (α, ý, ψ, and p120) and actin cytoskeleton, which plays a pivotal role in maintaining epithelial cell polarity and integrity (McCrea and Park, 2007). In our scRNA-seq study on EKP GOs, loss of Cdh1 resulted in transcriptional reprogramming and altered cell proportions, specifically reducing the Aqp5^high^ cluster and increasing the Mki67^high^ cluster (**Fig. 3J, K**). AQP5 is specifically expressed in pyloric stem cells, as well as being frequently expressed in gastric cancers and their metastases (Tan et al., 2020). Meanwhile, Aqp5 was expressed in a subpopulation of gastric cancer cells, some of which were KI67+. Our unbiased scRNA-seq analysis distinctly revealed two separate clusters, namely Aqp5^high^ cells and Mki67^high^ cells (**Fig. 3J**), denoting that the Aqp5^high^ cells within our murine GO model are not proliferative. This finding aligns with Barker’s study, wherein some Aqp5+ cells were found to be KI67 negative (Tan et al., 2020). Remarkably, in EKP organoids, we observed a reduction in Aqp5^high^ cells alongside an increase in Mki67^high^ cells (**Fig. 3J, K**). Consistently, in EKP tumors, a higher proportion of proliferative cells was observed compared to KP tumors (**Fig. 4G, I**). These outcomes suggest that Mki67^high^ cells might represent cells-of-origin in EKP tumors, characterizing the DGAC1 subtype with CDH1 loss. However, further rigorous experiments are warranted to validate this observation.

Intriguingly, CDH1 loss activates EZH2 regulon, and EZH2 blockade suppresses EKP tumor growth (**Fig. 5**). EZH2 modulates gene expression in various ways: gene repression via Polycomb Repressive Complex 2 (PRC2)-dependent histone methylation, PRC2-dependent non-histone protein methylation, or gene activation via transcriptional activator complex. The detailed mechanisms of how EZH2 is engaged in CDH1 loss-associated DGAC tumorigenesis remain to be determined. Nonetheless, given that an EZH2 inhibitor (tazemetostat) is clinically available, targeting EZH2 would be a viable option for the DGAC1 subtype in addition to T cell-based ICIs. The use of epigenetic modulators has been found to enhance the infiltration of effector T cells, suppress tumor progression, and improve the therapeutic effectiveness of PD-L1 checkpoint blockade in prostate or head and neck cancer (Jadhav et al., 2019; Weber et al., 2021). Additionally, pharmacological inhibition of EZH2 has been shown to inhibit tumor growth and enhance the efficacy of anti-CTLA-4 treatment in bladder cancer (Wherry and Kurachi, 2015). Given the enriched expression of immune checkpoints in DGAC1 (**Fig. 2J-L**), a combination therapy involving EZH2 inhibitors and ICIs may hold potential benefits for DGAC1 patients. Moreover, it should be determined whether EZH2-induced transcriptional reprogramming mediates CDH1 loss-induced transcriptional reprogramming of DGAC1.

Another remaining question is how CDH1 loss activates the EZH2 regulon. Mesenchymal cells re-wire PI3K/AKT signaling to stimulate cell proliferation (Salt et al., 2014). Additionally, it was shown that PI3K/AKT signaling is required for EZH2 activity in *KRAS^G12D^* mutant cells (Riquelme et al., 2016). Thus, it is plausible that EMT-activated PI3K/AKT signaling might activate EZH2. Consistent with this, compared to DGAC2, the DGAC1 subtype shows high scores for EMT and PI3K/AKT/MTOR pathways, and low score for inversely related EZH2 downstream target gene expression (**fig. S1J, S1M, Fig. 5I, J**).

Limitations of scRNA-seq include relatively shallow sequencing depth and restricted information not overcoming intra-tumoral heterogeneity. Thus, increasing the number of scRNA-seq datasets and spatial transcriptomics should follow in future studies. Furthermore, although this is the first stratification of DGAC by single-cell transcriptome, the pathological relevance of CDH1 status (or alternative molecular signatures; **Fig. 1M**) with ICI response remains to be clinically demonstrated.

Together, our study stratifies DGAC patients by integrative single-cell transcriptomics with experimental validation and unravels an unexpected role of CDH1 in restricting transcriptional reprogramming and immune evasion of DGAC, which provides new insight into the biology of DGAC tumorigenesis and helps improve immunotherapy efficacy.

## Author contributions

G.Z., Y.H., and J.-I.P. conceived and designed the experiments. G.Z., Y.H., S.Z., K.-P.K., B.K., J.Z., S.J., and V.V. performed the experiments. G.Z., Y.H., S.Z., K.-P.K., B.K., S.S., J.A.A., and J.-I.P. analyzed the data. M.P.P., Y.F., S.S., and J.A.A. provided the sequencing files and clinical data for human scRNA-seq analyses. N.N. and H.W. read and analyzed the stained slides. G.Z., Y.H., S.Z., K.-P.K., B.K., and J.-I.P. wrote the manuscript.

## Acknowledgments

We are grateful to Pierre D. McCrea, Malgorzata Kloc, Rachael Miller, and Adriana Paulucci for their insightful comments. This work was supported by the Cancer Prevention and Research Institute of Texas (RP200315 to J.-I.P.). The core facilities at MD Anderson (DNA Sequencing and Genetically Engineered Mouse Facility) were supported by National Cancer Institute Cancer Center Support Grant (P30 CA016672). This work was performed at the Single Cell Genomics Core at BCM partially supported by NIH-shared instrument grants (S10OD023469, S10OD025240) and P30EY002520.

## Methods

### Mice

All mouse experiments were approved by the MD Anderson Institutional Animal Care and Use Committee and performed under MD Anderson guidelines and the Association for Assessment and Accreditation of Laboratory Animal Care international standards. Compound transgenic mice *Kras^LSL-G12D/+^*; *Trp53^fl/fl^* (KP) mice have been previously described (Kim et al., 2021). C57BL/6 mice were purchased from the Jackson Laboratory (Maine, USA).

### Gastric organoids generation

The protocol for generating gastric organoids (GOs) was previously described (Bartfeld et al., 2015). The mice were sacrificed, and the mouse stomach was collected, and the forestomach was removed. Then, the reserved stomach tissue was cut through the lesser curvature, and the stomach was rinsed with ice-cold PBS with 1% penicillin/streptomycin to remove blood. The tissue samples were carefully immersed in chelating buffer (sterile distilled water with 5.6 mmol/L Na_2_HPO_4_, 8.0 mmol/L KH_2_PO_4_, 96.2 mmol/L NaCl, 1.6 mmol/L KCl, 43.4 mmol/L sucrose, 54.9 mmol/L D-sorbitol, 0.5 mmol/L DL-dithiothreitol, pH 7) in a 10 cm dish, then the tissue was transferred to a dry dish. The epithelial layer was peeled and minced into pieces using forceps. Minced epithelial pieces were placed into 10 mL cold chelating buffer, followed by robust pipetting up and down to rinse the tissue until the supernatant was clear. A 20 mL chelating buffer was prepared with 10 mM EDTA under room temperature, and the tissue was incubated in there for 10 min. The tissue was tenderly pipetted gently once up and down, and the pieces were allowed to settle. The tissue was then moved to the clean bench. Most of the water was removed, and the tissue pieces were carefully placed in the middle of a sterile 10 cm dish. A glass microscopy slide was put on top of the tissue and pressure was added upon the slide until the tissue pieces seemed cloudy. The cloudy tissue pieces were then flushed from the slides in 30 mL of cold Advanced DMEM/F12. The large tissue fragments were allowed to sediment by gravity. The cloudy supernatant was transferred to two 15 ml tubes. The tubes were then centrifuged for 5 min at 200 g and 4°C. The supernatant was carefully removed and resuspended with Matrigel-medium mixture (12 μL Matrigel mix with 8 μL GOs culture medium/well). Approximately 40 glands per 20 μL Matrigel-medium mixture per well of a 48-well plate were seeded. The plate was steadily transferred to the incubator to let it solidify for 10 minutes. Then, 500 μL of GOs culture medium was added to cover the dome, and the plate was incubated at 37 °C with 5% CO2. The medium was changed every 2 days.

### Gastric organoids culture

**Table S1** was referred to for the culture medium ingredient. The organoids were passaged using the following steps: **1.** The culture medium was discarded. **2.** The Matrigel was scraped with a pipette tip and dissociated by pipetting. **3.** The organoids were collected from three wells (48-well) in the 15 mL tube with cold medium. **4.** The supernatant was discarded after centrifugation at 1000 RPM and 4°C. **5.** The dissociated organoids were washed with 13 mL of cold 1× PBS, centrifuged (1000 RPM, 4 min), and the supernatant was removed. **6.** The organoids were resuspended in 1 mL of Trypsin-EDTA (0.05%). **7.** The sample was transferred to a 1.7 mL Eppendorf tube, then pipetted up and down. **8.** The sample was incubated in a 37 °C with 5% CO2 incubator for 30 min to 45 min. **9.** The tube was vibrated every 10 min. **10.** The organoid structure was further broken down by pipetting up and down. **11.** The sample was checked under the microscopy to ensure the organoids digested into cells. **12.** The sample was passed through the 35 μm cell strainer. **13.** The Trypsin was inactivated with 10% FBS medium and pipetted vigorously. **14.** The sample was collected in the 15 mL tube and centrifuged for 4 min at 1000 RPM. **15.** The supernatant was aspirated, and the cells were resuspended with GOs culture medium. **16.** The cells were counted, viability was checked, and the appropriate number of cells was calculated. **17.** Every 8 μL of cell suspension was mixed with 12 μL of Matrigel as a mixture and seeded in the 48-well plate. **18.** The plate was transferred to the incubator and allowed to solidify for 10 minutes. **19.** 500 μL of GOs culture medium was added to cover the dome and incubated at 37 °C with 5% CO2. **20.** The medium was changed every 2 days.

The organoids were cryopreserved as follows: The organoids were dissociated following above **organoid passaging (step1-15)** protocol. The cells were then added with 10% volume of DMSO and transferred to the cryovials.

### CRISPR/Cas9-based gene knockout in GOs

Knockout (KO) of *Cdh1* was performed by CRISPR/Cas9 genome editing using pLentiCRISPRv2 (Addgene plasmid #52961) according to Zhang laboratory’s protocol (Ran et al., 2013). Five single guide RNA (sgRNA) targeting *Cdh1* were designed using CRISPick (https://portals.broadinstitute.org/gppx/crispick/public) and cloned into a pLentiCRISPRv2-puro vector. An empty sgRNA vector was used as a negative control. The five targeting sequences against *Cdh1* were: #1: 5’-ATGAT GAAAA CGCCA ACGGG-3’, #2: 5’-ACCCC CAAGT ACGTA CGCGG-3’, #3: 5’-TTACC CTACA TACAC TCTGG-3’, #4: 5’-AGGGA CAAGA GACCC CTCAA-3’, and #5: 5’-CCCTC CAAAT CCGAT ACCTG-3’. sgRNA 1# (5’-ATGAT GAAAA CGCCA ACGGG-3’) was successfully knock out *Cdh1* in GOs. See **Table S2** for primer sequence to validate *Cdh1* knockout efficiency.

### Lentivirus production and transduction

The HEK293T cells were co-transfected with 5 μg of constructs, 5 μg of plasmid Δ8.2 (Plasmid #8455, Addgene), and 3 μg of plasmid VSVG (Plasmid #8454, Addgene) in a 10 cm dish. The cells were incubated at 37°C, and the medium was replaced after 12 h. The virus-containing medium was collected 48 h after transfection. The organoids were dissociated following the **organoid passaging protocol (step 1-14)**, and the supernatant was aspirated, leaving the pellet. For transduction, 20 μL of cell suspension was used. The amount of polybrene (8 μg/mL) was calculated and mixed with virus-containing medium before adding to the cells. The polybrene containing virus medium was added to the cell pellet, and the cell suspension was transferred to a 1.7 mL Eppendorf Tube. The tube was centrifuged at 600 g at 37 °C for 1 h. Without disturbing the cell pellet, the tube was incubated in the 37 °C incubator for 4 h. The supernatant was then removed, and the cell pellet was resuspended with the required volume of GOs culture medium (8 μL for one well of 48-well plate) and placed on ice for cool down. The appropriate volume of pre-thawed Matrigel (12 μL for one well of 48-well plate) was added to the tube, and the dome was seeded in the middle of a 48-well plate. The plate was then incubated for 10 min at 37 °C with 5% CO2. GOs culture medium was added to the well. After 48 h, the infected organoids were selected with 1 μg/mL puromycin.

### Adenovirus transduction

We used Adeno-Cre virus to treat *Kras^LSL-G12D/+^*; *Trp53^fl/fl^* organoids. The protocol was previously described (Ko et al., 2022; Ko et al., 2023). The cells were first dissociated from GOs as described in the **organoid passaging protocol (step 1-14)**. The cell number was counted, and the ratio of adenovirus: organoid cell was 1000 PFU/μL:1 cell. The cell suspension, virus-containing medium, and Matrigel were mixed, and the drop was placed in the center of the well. The cell suspension and virus-containing medium were mixed before adding GOs culture medium up to 8 μL. Then, 12 μL of Matrigel was added to the mixture on ice. The plate was incubated in the 37°C cell culture incubator for 15 min to allow the Matrigel to solidify. After 48 h, the infected organoids were treated with 10 μM Nutlin-3 to select *Trp53* KO organoids. The primer sequence to validate *Trp53* KO and *Kras^G12D/+^* can be found in **Table S2**.

### Organoid imaging and size measurement

After 7 days of organoid seeding in Matrigel, the size of the organoids was analyzed by measuring the volume under the microscope (ZEN software, ZEISS). To reduce the vulnerability of GOs, the measurements were conducted more than 3 passages after isolation from the knockout experiments. All experiments included more than 50 organoids per group.

### Tissue microarray

DGAC cancer tissue microarray slides contained 114 patients’ samples. Patients’ information is shown in **Table S4**.

### Histology and immunohistochemistry

All staining was performed as previously described (Jung et al., 2018). For organoids staining, 7 days after seeding, GOs were collected by dissociating Matrigel mixture using ice-cold PBS and fixed in 4% paraformaldehyde at room temperature. For tumor tissue, excised tumors were washed with ice-cold PBS and fixed with formaldehyde at room temperature. After paraffin embedding, tumor tissue and organoid sections were mounted on microscope slides. For H&E staining, sections were incubated in hematoxylin for 3-5 min and eosin for 20-40 s. After washing with tap water, slides were dehydrated, and the coverslips were mounted with mounting media. For immunofluorescence staining, after blocking with 5% goat serum in PBS for 1 hr at room temperature, sections were incubated with primary antibodies (MKI67 [1:200], CDH1 [1:200], CD3 [1:200], CD8 [1:200], CD4 [1:200], PDCD1 [1:200], TIM3 [1:200], CD11B [1:200], LY6G [1:200]) overnight at 4 °C and secondary antibody (1:250) for 1 hr at room temperature in dark. Sections were mounted with ProLong Gold antifade reagent with DAPI (Invitrogen). For immunohistochemistry staining, after blocking with 5% goat serum in PBS for 1 hr at room temperature, sections were incubated with primary antibodies (CDH1 [1:200], H3K27Me3 [1:200], H3K27Ac [1:200]) overnight at 4 °C and secondary antibody (1:250) for 1 hr at room temperature in dark. Incubate the slides in the DAB solution until tissue become brown and background still white. Observed under the microscope until the strongest signal shows and stop reaction with tap water wash. Used the same incubation time for same antibody on different slides. Sections were incubated in hematoxylin for 3-5 min and mounted with mounting media. Images were captured with the fluorescence microscope (Zeiss; AxioVision). See **Table S3** for antibody information.

### 2D culture

The organoids were dissociated following the **organoid passaging protocol (step1-14)**. The supernatant was aspirated and then resuspended with DMEM + 10% FBS with 10 μM Y-27632, and the organoids were seeded on a 24-well plate. Cells were passaged every 3-5 days. After the third passage, Y-27632 was removed from the culture medium. DMEM supplemented with 10% FBS and 10% DMSO was used to freeze cells and store them in liquid nitrogen.

### Allograft transplantation

Five-week-old C57BL/6 mice were maintained in the Division of Laboratory Animal Resources facility at MD Anderson. 2D-cultured KP and EKP cells (1 × 10^6^) were injected subcutaneously into both flanks of mice. Tumor volume was calculated by measuring with calipers every 3-4 days (volume = (length × width^2^)/2). Mice were euthanized, and tumors were collected at day 15. The excised tumors were photographed and paraffin-embedded for immunostaining. For GSK343 treatment, 2D-cultured EKP cells (1 × 10^6^) were injected subcutaneously into both flanks of mice. After the tumors were palpable, we performed the first measurement with calipers. We divided the mice into two groups of three mice each and administered DMSO and GSK343 (20 mg/kg) intraperitoneally every other day. The initial tumor volumes between the two groups were comparable. Tumor volume was calculated by measuring with calipers every 3-4 days (volume = (length × width^2^)/2). Mice were euthanized, and tumors were collected at day 20.

### Crystal violet staining

Cells (1 × 10^3^) were seeded on a 6-well plates, and the medium was replaced every 2 days. Plates were rinsed with 1× PBS, fixed with 4% paraformaldehyde solution for 20 min, and stained with crystal violet solution (0.1% crystal violet, 10% methanol) for 20 min, followed by rinsing with tap water.

### Gastric organoids library preparation for scRNA-seq

For scRNA-seq, organoids from WT, KP, and EKP were collected 7 days after seeding and follow the **organoid passaging (step1-14)** protocol. After trypsin had been inactivated with 10% FBS DMEM, a single-cell suspension was collected by passing cells through a 70 μm cell strainer and followed by a 40 μm cell strainer. Each group was tagged with two CMO tags from the CellPlex kit (10ξ Genomics). The tagged cells of each group were pooled together with the same number of cells after being counted. Single cell Gene Expression Library was prepared according to Chromium Single Cell Gene Expression 3v3.1 kit with Feature Barcode technology for cell Multiplexing (10x Genomics). In brief, tagged single cells, reverse transcription (RT) reagents, Gel Beads containing barcoded oligonucleotides, and oil were loaded on a Chromium controller (10x Genomics) to generate single cell GEMS (Gel Beads-In-Emulsions). Incubation of the GEM produced barcoded, full-length cDNA as well as barcoded DNA from the cell Multiplexing. Subsequently the GEMS are broken and pooled. Following cleanup using Dynabeads MyOne Silane Beads, full-length cDNA is amplified by PCR for library prep through fragmentation, end-repair, A-tailing, adaptor ligation and amplification, while the barcoded DNA from the cell Multiplexing is amplified for library prep via PCR to add sequencing primers. The cDNA library was sequenced on an Illumina NovaSeq platform (Novogene), mapped to the GRCm38/mm10 genome, and demultiplexed using CellRanger. The resulting count matrices files were analyzed in R (Seurat) or Python (Scanpy).

### scRNA-seq - raw data processing, clustering, and annotation

We used Cell Ranger to perform demultiplexing and reads alignment of sequencing raw data for the scRNA-seq matrices generation. Ambient RNA and doublets were removed by SoupX (Young and Behjati, 2020) and Scrublet (Wolock et al., 2019), respectively. Scanpy(Wolf et al., 2018) was used for processing the scRNA-seq data. For the organoid dataset, cells with less than 50 genes expressed and more than 30% mitochondrial reads, 30% rpl reads, and 25% rps reads were removed. Genes expressed in less than 5 cells were removed. Then we normalized and log-transformed the gene expression for each cell. The percentages of mitochondrial reads, rpl reads, and rps reads were regressed before scaling the data. We reduced dimensionality and cluster the cells by Leiden (resolution=0.5). Cell lineages were annotated based on algorithmically defined marker gene expression for each cluster (sc.tl.rank_genes_groups, method=‘t-test’). See **Table S17**, the most differentially expressed 100 genes of each cluster were listed. For the DGAC dataset, cells with less than 100 genes expressed and more than 80% mitochondrial reads, 30% rpl reads, and 25% rps reads were removed. Genes expressed in less than 25 cells were removed. Normalization, log-transformation, regression, dimensionality reduction, and Leiden clustering (resolution=1) were the same as the way we use in organoids. Cell lineages were annotated based on algorithmically defined marker gene expression for each cluster (sc.tl.rank_genes_groups, method=‘t-test’). See **Table S6, S7, and S8** for details, the most differentially expressed 100 genes of each cluster or type were listed. For the DGAC dataset merged with normal stomach dataset, cells with less than 100 genes expressed and more than 100% mitochondrial reads, 40% rpl reads, and 30% rps reads were removed. Genes expressed in less than 25 cells were removed. Normalization, log-transformation, regression, dimensionality reduction, and Leiden clustering (resolution=1) were the same as the way we use in organoids. Cell lineages were annotated based on algorithmically defined marker gene expression for each cluster (sc.tl.rank_genes_groups, method=‘t-test’). See **Table S10** for details, the most differentially expressed 100 genes of each cluster were listed. More information about the software and algorithms used in this study is shown in **Table S19**.

### Proportion difference analysis

The cell number of each cluster were retrieved by Scanpy (adata.obs[’leiden’].value_counts()). We analyzed and plotted the differences between clusters from the two datasets using the GraphPad Prism 9.4. Then we grouped each cell cluster from the integrated dataset and compared the cluster differences between the two datasets.

### Regulon analysis

For the gene regulatory network inference in organoids, we used the pySCENIC package (Van de Sande et al., 2020) to compute the specific regulons for each cell cluster. The Loom file of each organoid dataset was used, and the regulon pattern-based UMAP was redrawn based on the AUCell scoring method (Aibar et al., 2017). Regulon specificity score (RSS) (Suo et al., 2018) and Z score were used to determine how specific the regulon is for one certain cell cluster. More specific the regulon is, the higher RSS or Z score is for one certain cluster. Following the criteria that RSS and Z score should be high at the same time, we identified 20 regulons that specific to EKP. These processes were repeated five times in each organoid dataset (WT, KP, and EKP). RSS of regulons from each mouse gastric organoid dataset (WT, KP, and EKP) was listed in **Table S18**.

### Scissor analysis

To determine the pathology of murine organoids, we compared the transcriptomic similarity of the organoids scRNA-seq dataset and the bulk RNA-seq datasets of DGAC patients by Scissor package (Sun et al., 2022). The RNA-seq data of tumor and the adjacent normal samples of DGAC patients were downloaded from the GDC data portal (TCGA-STAD). The murine genes were converted to human homologs by biomaRt. The Scissor analysis was performed by using the Cox regression model (alpha = 0.32). The goal of Scissor is to identify a small group of cells that are most highly correlated with the specific phenotypes with high confidence. Based on this motivation as a priori, we determined *α* using the following criteria: the number of Scissor-selected cells should not exceed a certain percentage of total cells (default 20%) in the single-cell data (Sun et al., 2022).

### Cell-cell communication analysis

‘CellChat’ (Jin et al., 2021) package in R (https://www.r-project.org) was used to analysis the ligand-receptor interaction-based cell-cell communication in scRNA-seq datasets. The integrated dataset was processed, clustered, and annotated using the scanpy package (Wolf et al., 2018) in python, then transformed into .rds files. Transformed datasets were analyzed by CellChat with default parameters (p-value threshold = 0.05).

### Pathway score analysis

Pathway score was analyzed by Scanpy (Wolf et al., 2018) with the ‘scanpy.tl.score_genes’ function (Wolf et al., 2018). The analysis was performed with default parameters and the reference genes from the gene ontology biological process or the Kyoto Encyclopedia of Genes and Genomes database (Kanehisa, 1996; Ashburner et al., 2000). The gene list for the score analysis is shown in **Table S9**.

### Human scRNA-seq data analysis

The scRNA-seq data set of 19 DGAC patients’ samples (Patients information is shown in **Table S5**) has been previous reported from our group and the detailed clinical and histopathological characteristics are described (EGAS00001004443) (Wang et al., 2021). The meta data of the scRNAseq is presented on **Table S5.**

The scRNA-seq data set of the 29 normal adjacent stomachs (GSE150290) (Kim et al., 2022) was extracted from the Gene Expression Omnibus (GEO) database and analyzed with Scanpy and Python (Wolf et al., 2018). The 19 DGAC patients’ datasets were integrated and clustered by Scanpy (Wolf et al., 2018) for the subclassification of DGACs based on CDH1 inactivation. The 19 DGAC patients’ datasets and 29 normal adjacent stomachs were integrated and clustered in Scanpy (Wolf et al., 2018) for later infercnvpy analysis. “Harmony” (Korsunsky et al., 2019) algorithm was used to remove batch effects. Then, the dendrogram and correlation matrix heatmap were plotted with Scanpy (Wolf et al., 2018). The dendrogram shows the distance of each dataset based on principal component analysis, and the correlation matrix heatmap shows Pearson correlation by a color spectrum.

### Copy number variation analysis

To detect the genomic stability of groups DGAC1, DGAC2, we performed copy number variations (CNVs) inference from the gene expression data using the Python package infercnvpy (https://icbi-lab.github.io/infercnvpy/index.html). We performed infercnvpy on DGAC1, DGAC2 using the Normal group (29 human normal adjacent stomachs) as reference. The gene ordering file which is containing the chromosomal start and end position for each gene was created from the human GRCh38 assembly. The GRCh38 genomic positions annotated file was downloaded from https://support.10xgenomics.com/single-cell-gene-expression/software/downloads/latest. Infercnvpy was used to plot chromosome heatmap and CNV scores in the UMAP.

### Gene set enrichment analysis (GSEA)

GSEA was conducted via the R package “fgsea” (Korotkevich et al., 2021) according to the DEG list generated by Scanpy. NES (Normalized Enrichment Score) represents the degree of enrichment of a gene set in a given dataset, measuring the coordinated upregulation or downregulation of genes within the set compared to a reference condition. It is normalized to account for variations in gene set size and dataset characteristics, providing a more robust measure of enrichment. The enrichment value was calculated and plotted with the fgsea package (permutation number = 2,000). All enriched pathways were listed in **Table S11-S16**.

### Public sequencing database

All TCGA cancer patients’ sequencing data referenced in this study were obtained from the TCGA database at cBioPortal Cancer Genomics (http://www.cbioportal.org).

## Data availability

scRNA-seq data are available via the GEO database (GSE226266; log-in token for reviewers: edazwaukzvsxbop).

## Code availability

The code used to reproduce the analyses described in this manuscript can be accessed via GitHub (https://github.com/jaeilparklab/EKP_DGAC_project) and will also be available upon request.

## Statistical analyses

GraphPad Prism 9.4 (Dogmatics) was used for statistical analyses. The Student’s *t*-test was used to compare two samples. The one-way ANOVA was used to compare multiple samples*. P* values < 0.05 were considered statistically significant. Error bars indicate the standard deviation (s.d.) otherwise described in figure legends.

## Supplementary Figures

**Supplementary Figure S1.**
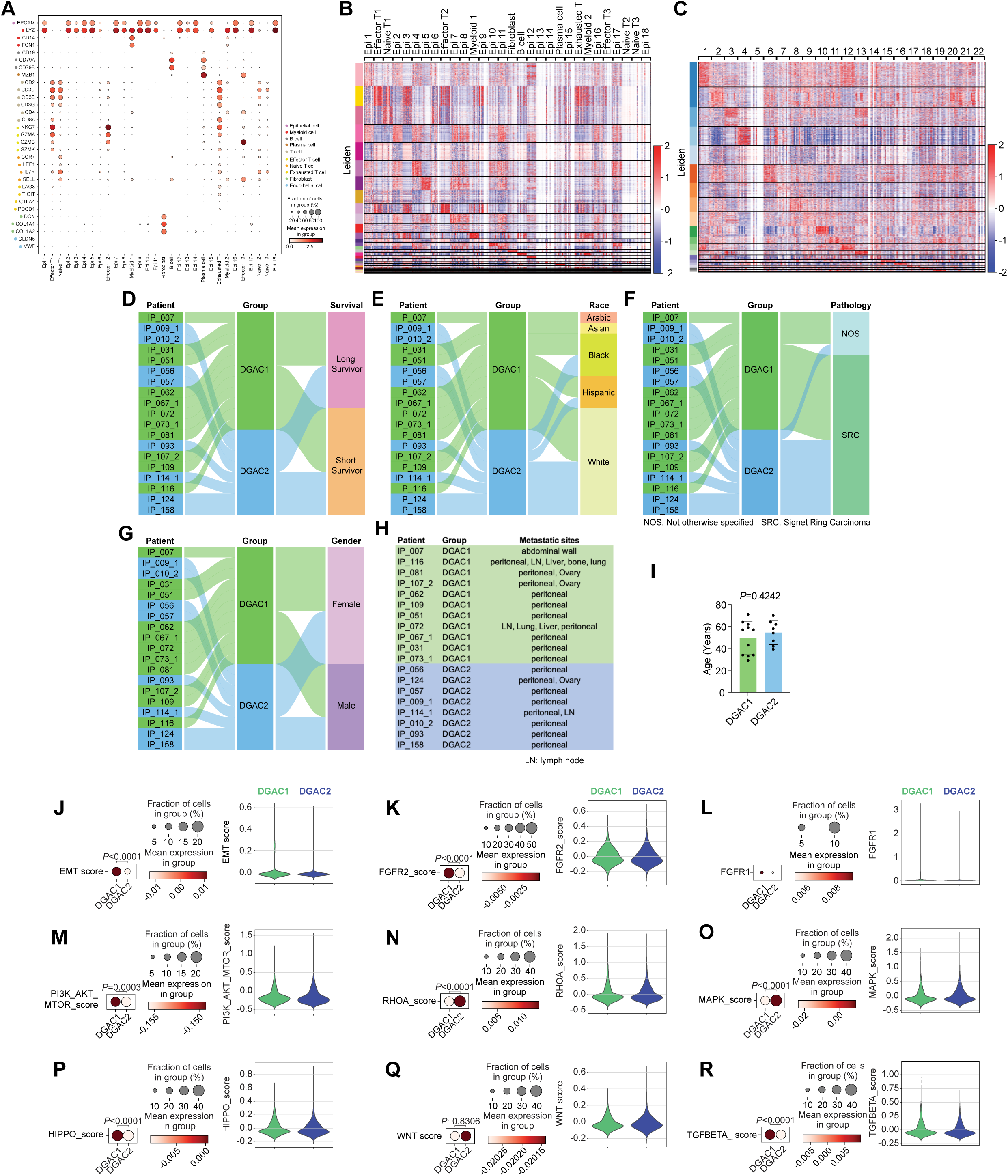
Transcriptional, clinical, and molecular characterization of DGAC subtypes. **A.** Dot plots of epithelial cell, myeloid cell, B cell, plasma cell, T cell, effector T cell, naïve T cell, exhausted T cell, fibroblast, and endothelial cell markers in merged 19 DGAC patients scRNA-seq data. **B.** Leiden-based heatmap of all cells of merged datasets with annotation in 19 DGAC patients. The most highly variable 100 genes of each cluster were showed in Table S6. **C.** Leiden-based heatmap of epithelial cells of merged datasets in 19 DGAC patients. The most highly variable 100 genes of each cluster were showed in Table S7. **D-G.** Venn diagram illustrating 19 DGAC patient groups with survival, race, pathology, and gender data. Long-term survivors (surviving over 1-year post-diagnosis) and short-term survivors (deceased within 6 months post-diagnosis) were classified according to our previous publication (Wang et al., 2021). **H.** Metastatic sites of DGAC1 and DGAC2 patients. **I.** Age difference between DGAC1 and DGAC2 patients. **J-R.** Dot plots, violin plots, and feature plots of EMT (J), FGFR2 (K), FGFR1 (L), PI3K_AKT_MTOR (M), RHOA (N), MAPK (O), HIPPO (P), WNT (Q), and TGFBETA (R) scores in two DGAC types. *P* values were calculated by using a *t*-test. The genes included in each score are listed in Table S9.

**Supplementary Figure S2.**
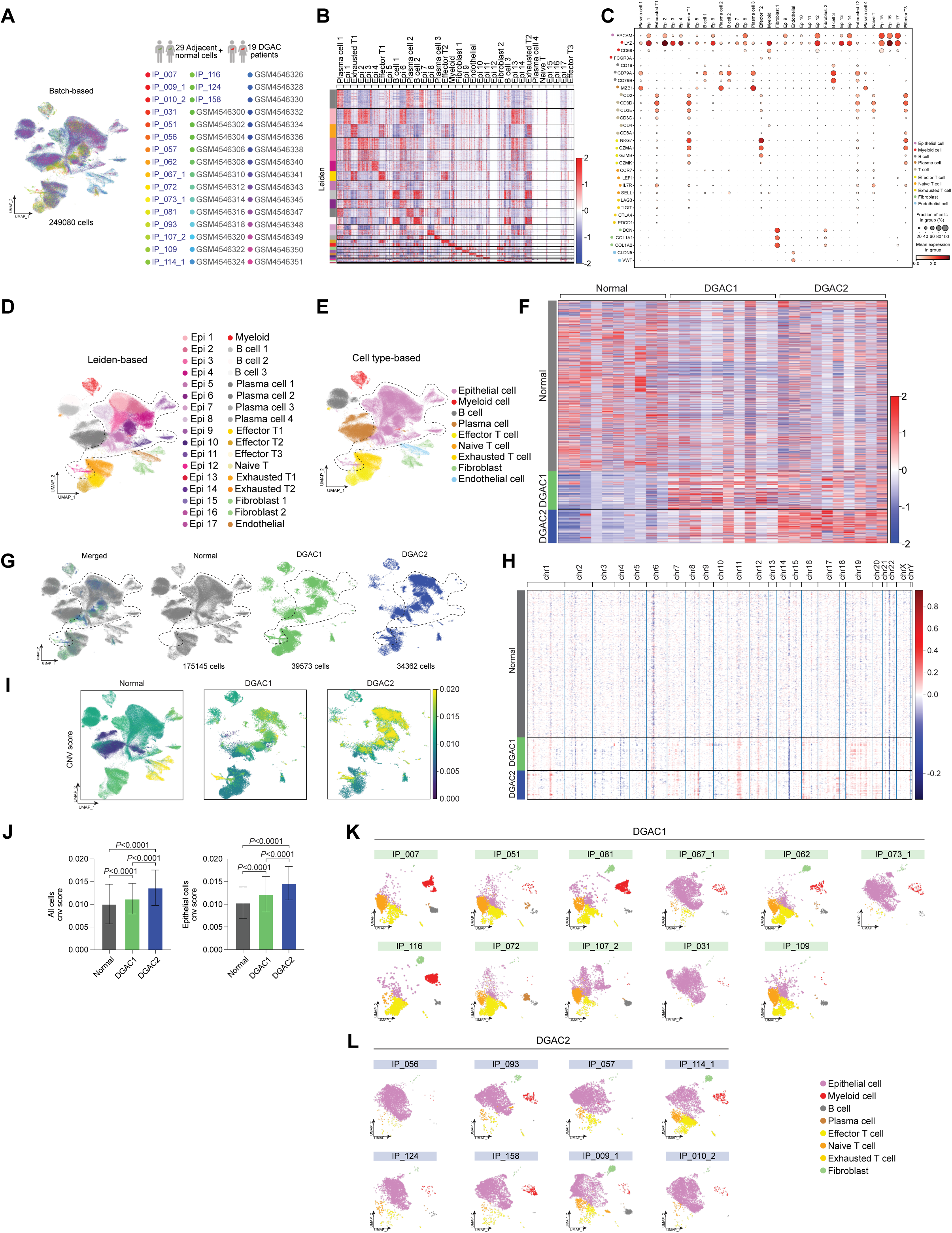
scRNA-seq analysis of 19 DGAC patients and 29 adjacent normal stomach tissue. **A.** Merged batch-based UMAP of 29 adjacent normal stomach tissue (Normal tissue) and 19 DGAC patients. Total cell numbers are 249080. Integration package: Harmony. **B.** Annotated Leiden-based integrated UMAPs of 19 DGAC patients and 29 adjacent normal stomach tissue. Epi: Epithelial cells; Myeloid: myeloid cells; Effector T: effector T cells; Naïve T: Naïve T cells; Exhausted T: Exhausted T cells; Endothelial: Endothelial cells. **C.** Dot plots of epithelial cell, myeloid cell, B cell, plasma cell, T cell, effector T cell, naïve T cell, exhausted T cell, fibroblast, and endothelial cell markers in merged 19 DGAC patients and 29 adjacent normal stomach tissue scRNA-seq data. **D.** Merged Leiden-based integrated UMAPs of 29 adjacent normal stomach tissue (Normal tissue) and 19 DGAC patients. Epi: epithelial cells; Myeloid: myeloid cells; Effector T: effector T cells; Naïve T: naïve T cells; Exhausted T: exhausted T cells. The most highly variable 100 genes of each cluster were showed in Table S10. **E.** Merged cell type-based UMAP of 29 Normal tissue and 19 DGAC patients. All cells were re-clustered according to the Leiden clusters and gathered as mega clusters. Dashed line-circle: epithelial cells. **F.** Type-based heatmap of all cells of merged datasets in 19 DGAC patients and 29 adjacent normal stomach tissue. **G.** Separated UMAPs of Normal tissue and two types of DGACs. Dashed line-circle: epithelial cells. **H.** CNV heatmap of DGAC1 and DGAC2, tumor-adjacent normal stomach tissue (Normal) was used as reference for the CNV inference. Red: copy number gain (CNG); blue: copy number loss (CNL) **I.** CNV heatmap of DGAC1 and DGAC2, tumor-adjacent normal stomach tissue (Normal) was used as reference for the CNV inference. **J.** Statistics analysis of CNV score of all cells (left panel) and epithelial cells (right panel) among Normal, DGAC1, and DGAC2. *P* values were calculated using the one-way ANOVA; error bars: SD. **K, L.** Individual cell type-based UMAP of the patients in DGAC1 and DGAC2. DGAC1 patients were enriched with stromal cells, mainly T cells. DGAC2 patients were enriched with epithelial cells.

**Supplementary Figure S3.**
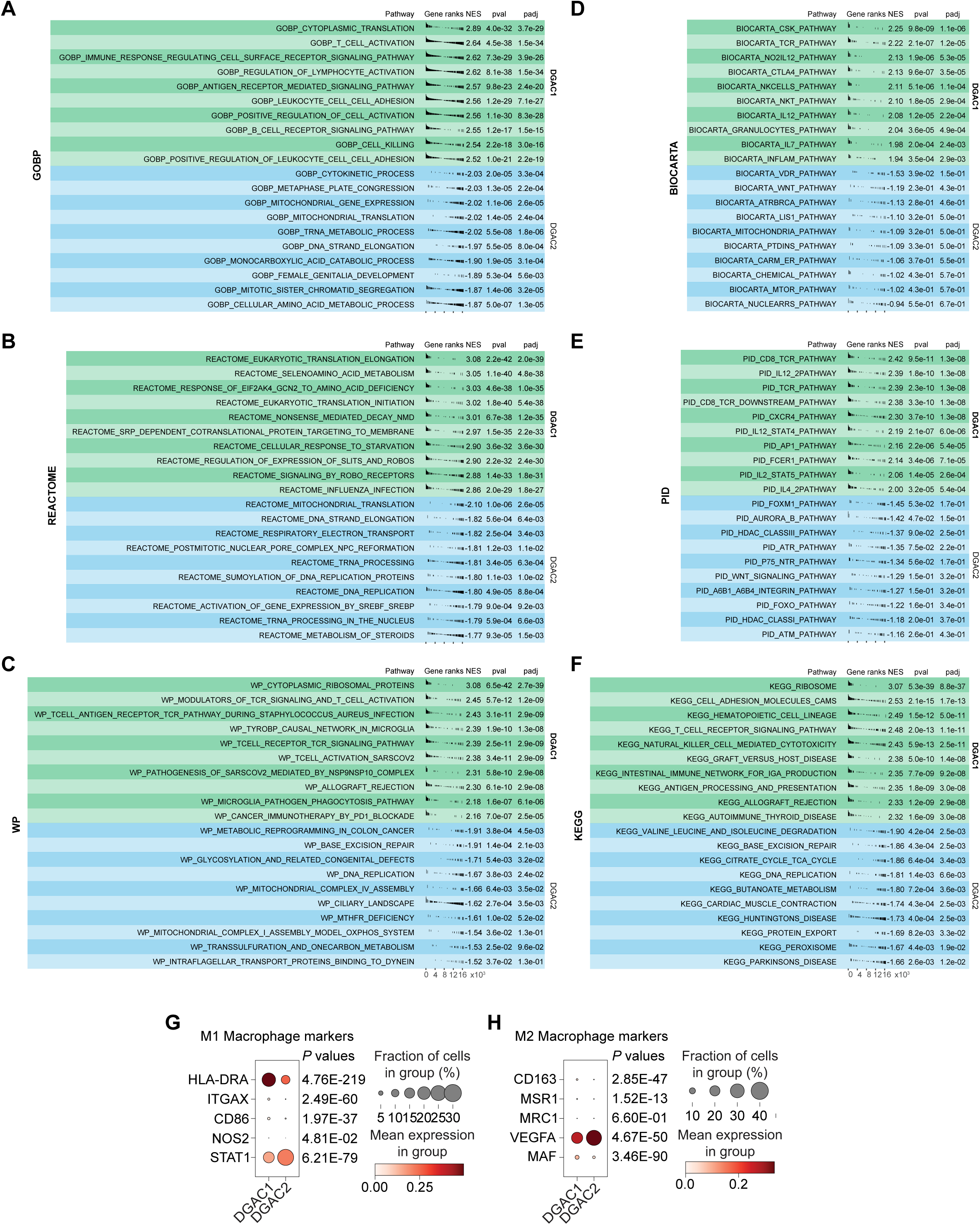
GSEA analysis and the expression of macrophage polarization markers of DGAC1 and DGAC2. **A-F.** GSEA analysis comparing DGAC1 to DGAC2 using DGAC2 as the reference gene set. Enriched pathways in DGAC1 are displayed in the upper green panel, while those in DGAC2 are shown in the lower blue panel. Pathway datasets analyzed include GOBP (A), REACTOME (B), WP (C), BIOCARTA (D), PID (E), and KEGG (F). Pathways with positive NES (Normalized Enrichment Score) indicate enrichment in DGAC1, while those with negative NES indicate enrichment in DGAC2. GOBP: Gene ontology biological process; REACTOME: Reactome gene sets; WP: WikiPathways gene sets; BIOCARTA: BioCarta gene sets; PID: PID gene sets; KEGG: KEGG gene sets. Pathways related with immune response were enriched in DGAC1 based on GOBP, WP, BIOCARTA, PID, and KEGG. **G, H.** Dot plot of macrophage polymerization markers in DGAC1 and DGAC2. Most of the M1 and M2 markers are enriched in DGAC1, except for STAT1 and VEGFA.

**Supplementary Figure S4.**
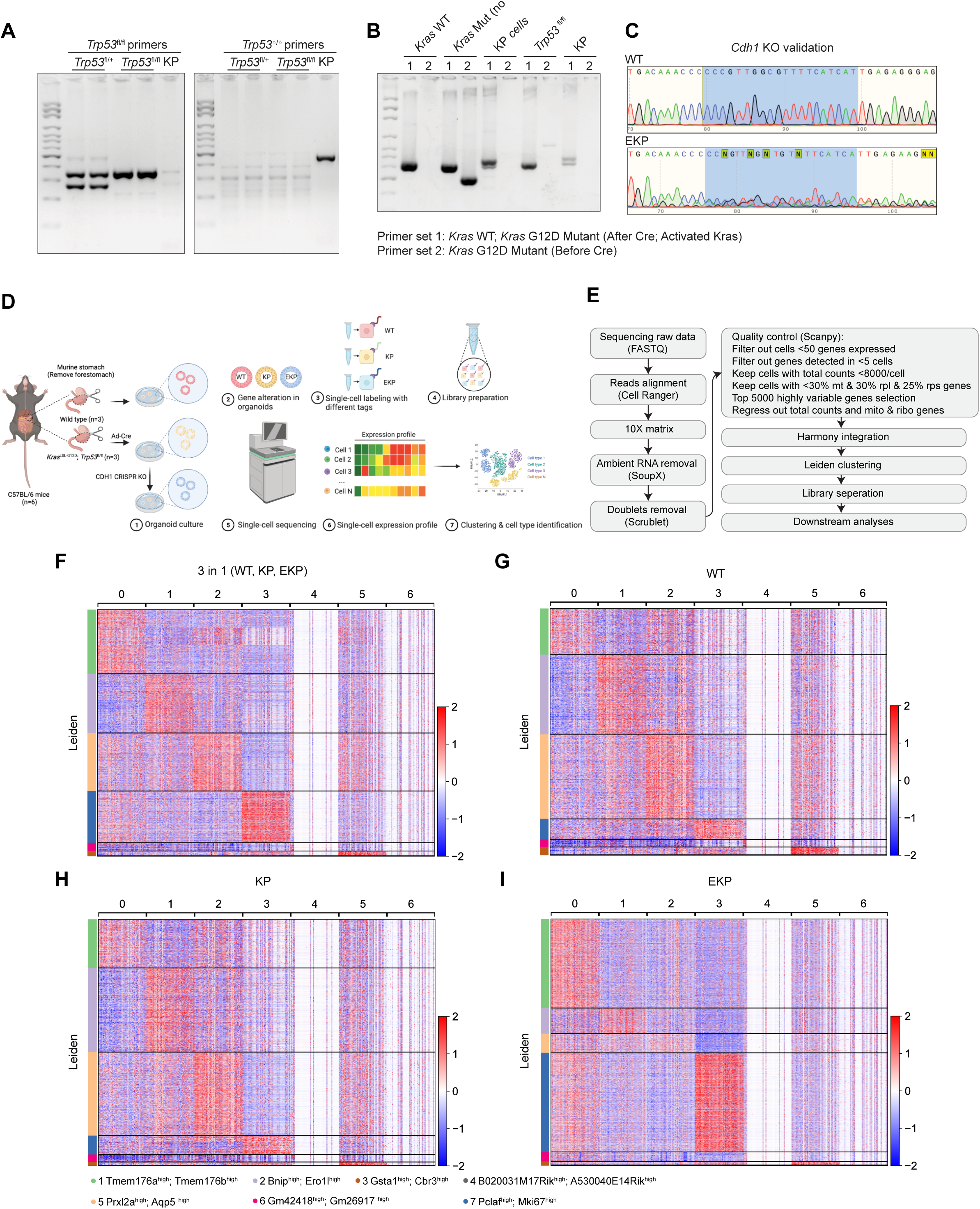
Validation of genetic engineering and scRNA-seq analysis of mouse GOs. **A-C.** Genotyping results of KP organoids (A and B). After adeno-Cre treatment, KP organoids lost *Trp53*, while *Kras^G12D^*was activated in KP organoids. After *Cdh1* CRISPR knock out (KO), we performed sanger sequencing to compare the sequence of *Cdh1* in WT and EKP (C). The five targeting sequences against *Cdh1* were showed in methods ‘**CRISPR/Cas9-based gene knockout in GOs**’. The primers used for genotyping were showed in Table S2. **D.** Illustration of the workflow for stomach tissue collection and dissociation, gene manipulation of the gastric organoids (GOs), sample preparation of multiplex scRNA sequencing. **E.** Workflow of single cell library preparation. **F.** Heatmap of each cell clusters of merged datasets, including WT, KP, and EKP. **G-I.** Separate heatmap of each cell clusters of WT, KP, and EKP datasets, respectively.

**Supplementary Figure S5.**
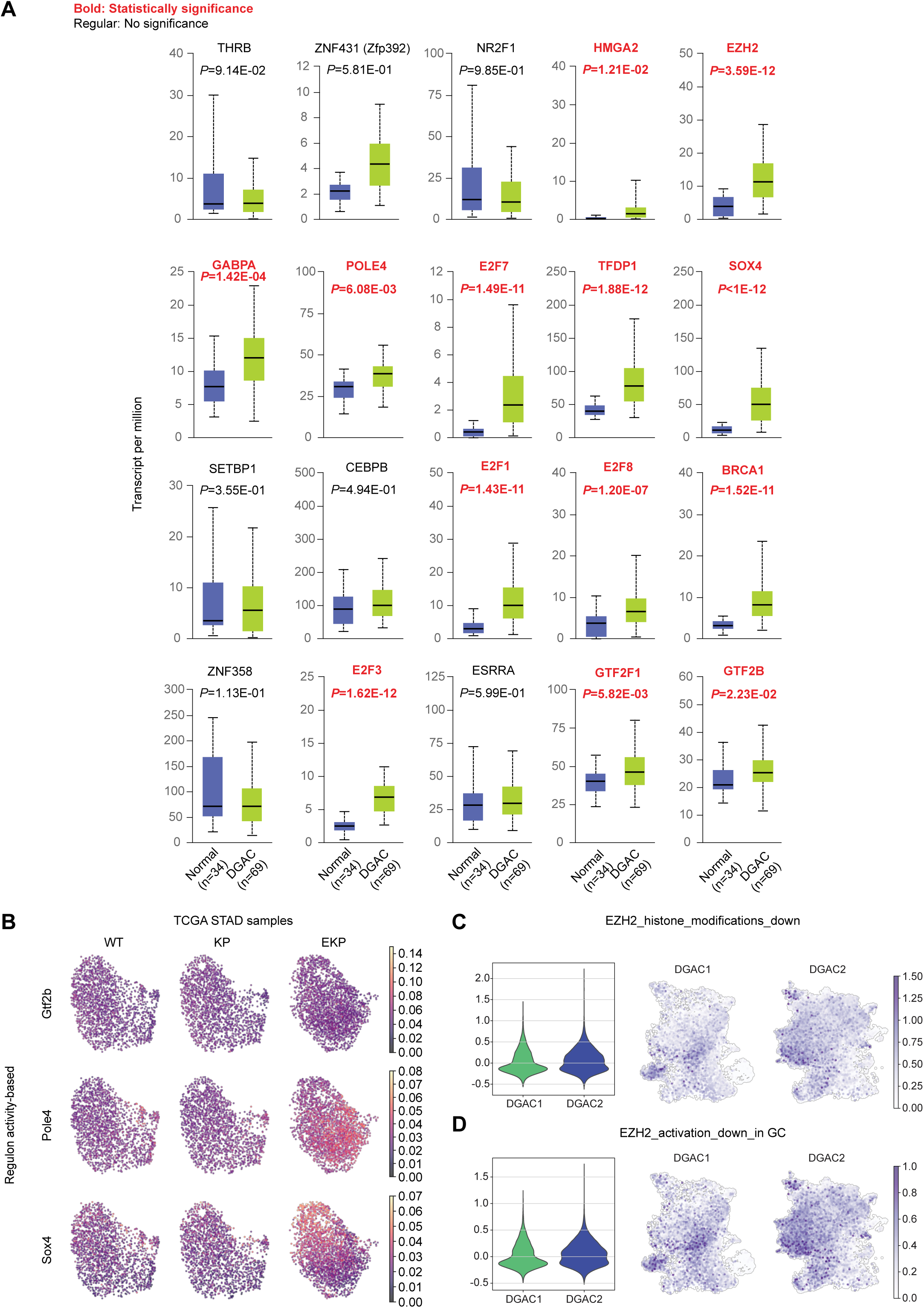
EKP-specific regulons expression and EZH2 downstream targeted genes expression. **A.** The expression of 20 regulons in TCGA DGAC patients and normal stomach. **B.** Regulon activity based UMAP of Gtf2b, Pole4, and Sox4. *P* values were calculated by using the Student’s *t*-test; error bars: SD. **C, D.** Violin (left panel) and feature plots (right panel) of EZH2 downstream target genes (C, genes which are downregulated by EZH2 through histone modification; D, genes which are downregulated by EZH2 reported in gastric cancer) scores in the epithelial cells of DGAC1 and DGAC2. Gene list of EZH2 targeted genes was listed in Table S9.

## Notes

### Competing Interest Statement

The authors have declared no competing interest.

### Summary of Updates

- New analysis of scRNA-seq datasets (19 instead of 20 patient tumors) - Clarification in scRNA-seq analysis - Additional experiments showing the impact of CDH1 loss on gastric organoid forming and differentiation.

